# Culture substrate stiffness impacts human myoblast contractility-dependent proliferation and nuclear envelope wrinkling

**DOI:** 10.1101/2023.08.08.552533

**Authors:** Jo Nguyen, Lu Wang, Wen Lei, Yechen Hu, Nitya Gulati, Carolina Chavez-Madero, Henry Ahn, Howard J. Ginsberg, Roman Krawetz, Matthias Brandt, Timo Betz, Penney M. Gilbert

## Abstract

The remarkable self-repair ability of skeletal muscle tissues is driven by muscle stem cells, whose activities are orchestrated by a variety of transient cues present in their microenvironment. Understanding how the cross-interactions of these biophysical and biochemical microenvironmental cues influences muscle stem cells and their progeny is crucial in strategizing remedies for pathological dysregulation of these cues in aging and disease. In this study, we investigated the cell-level influences of extracellular matrix ligands in the context of mechanically tuned culture substrates on primary human myoblast contractility and proliferation within 16 hours of plating. We found that fibronectin led to stronger stiffness-dependent responses in myoblasts compared to laminin and collagen as revealed by cell spreading area, focal adhesion morphology, traction stresses, and proliferation. While we did not observe stiffness-dependent changes in cell fate signatures at this early time-point, an analysis of the stiffness-sensitive myoblast proteome uncovered an enrichment for differentiation associated signatures on 17 kPa culture substrates, in addition to distinct profiles with relation to metabolism, cytoskeletal regulation, ECM interactions, and nuclear component enrichment. Interestingly, we found that softer substrates increased the incidence of myoblasts with a wrinkled nucleus, and that the extent of wrinkling could predict Ki67 expression. While nuclear wrinkling and Ki67 expression could be controlled by pharmacological manipulation of cellular contractility, cellular traction stresses did not correlate with nuclear envelope wrinkling level. These results provide new insights into the regulation of human myoblast stiffness-dependent contractility response by ECM ligands and also highlight a link between myoblast contractility and proliferation.

## Introduction

The human skeletal muscle system sustains body mobility for many years owing to its self-repair capacity, powered by muscle stem cells (MuSCs). These MuSCs are scattered throughout muscle tissues in between muscle fibers and exist mostly in a quiescence state during homeostasis. In the event of tissue damage, MuSCs become activated, begin to proliferate, and either fuse into muscle fibers or self-renew. These processes are carefully regulated by biophysical and biochemical cues at the MuSCs residence over the tissue repair period[1,2]. MuSCs reside in a niche that is structurally polarized with the basal lamina on one side and the muscle fiber sarcolemma on the other. The basal lamina is a sheet of extracellular matrix (ECM) composed of laminins, collagens, fibronectin, and proteoglycans. During tissue repair, the ECM surrounding MuSCs can be both produced and degraded by many cell types including MuSCs themselves, causing fluctuations in its composition and physical properties like stiffness[2]. Stiffening of bulk muscle tissue and single myofibers occur upon damage[3–5], aging[4,6–9], exercise[10], and disease[11–14], while engineered niche stiffness has been shown to control myogenic cell activation[15], proliferation[4,5,16–18], differentiation[4,5,13,16,17,19], self-renewal[20], and division symmetry[21]. However, less explored is the role of specific ECM ligands in regulating stiffness-sensing in myogenic cells. Since irregular ECM remodeling is often involved in skeletal muscle aging and disease pathology, it is necessary to extend our understanding of the synergy between the physical and compositional changes in the ECM in controlling myogenic cell behavior.

MuSC adhesion to the ECM is mediated by transmembrane integrin receptors, with the most well-known being integrin α7, who together with the β1 subunit form binding receptors to niche laminin. MuSCs can also express integrins α3[22] and α6[23,24] while MuSCs progenitors in vitro express more integrin subtypes[25]. Integrin subtypes are specific to the presented ECM ligands, therefore any compositional changes in the ECM immediate to the MuSCs microenvironment hypothetically would shift the profile of integrin subtypes recruited to the MuSCs membrane. For instance, laminin-α1 deposition has been reported to couple with the appearance of integrin α6 during MuSCs activation[24]. Since integrin subtypes can have differential binding mechanics with the ECM and thus tension-transmitting properties[26], ECM compositional dynamic not only could alter niche stiffness but potentially also how MuSCs “feel” that change. This hypothesis is challenging to test in the native MuSC niche, limited by the ability to manipulate physical niche factors without introducing chemicals disturbing the biochemical homeostasis. In vitro, murine MuSCs commitment to differentiation was reported to be sensitive to substrate stiffness on RGD-presenting hydrogels but not on laminin-presenting hydrogels[17]. MuSCs proliferation and migration have also been shown to be controlled by both ECM ligands and substrate stiffness by an unknown mechanism[17,27,28]. Deconstructing the interplay between ECM ligands and substrate stiffness converges at two questions: how ECM ligand identity changes the way myogenic cells sense substrate stiffness, and how substrate stiffness change ECM ligands signaling. In the current work, we contribute to the former question by characterizing the effects of ECM ligands on early adhesive and contractile response of myoblast to substrate stiffness, and then further emphasized the relationship between myoblast contractility and proliferation by reporting a correlation between nuclear envelope wrinkling level and Ki67 expression.

## Materials and Methods

### Fabrication of polyacrylamide (PAA) hydrogels

PAA hydrogels were fabricated according to the protocol described by Tse and Engler [29] with some modifications: hydrogels were cast on 35 mm glass bottom dishes (Matusunami Glass; D35-14-1-U) instead of coverslips, and deposition of NAOH was followed by immediate aspiration instead of heated evaporation. Hydrogels were then tethered with either fibronectin (Fibronectin bovine plasma, Sigma F1141), collagen (Collagen I, Rat Tail, Gibco A1048301), or laminin (Roche 11243217001) at a 100 mg/ml concentration according to Engler et. al. protocol. Expected Young’s moduli of the hydrogel formulations used in this study were extrapolated from the Engler et. al. protocol (i.e., 1.10±0.34, 2.55±0.17, 4.47±1.19, 8.44±0.82, and 16.70) and were then rounded to the nearest whole number: 1, 3, 5, 8, and 17 kPa, respectively, for simplicity.

For traction force microscopy, PAA hydrogels were fabricated following the same protocol above with the addition of embedding 100 µm fluorescent beads at a 1:200 dilution.

### Primary human myoblast preparation and expansion

Primary human myoblast cell lines were derived from human skeletal muscle biopsies. Four primary human myoblast cell lines were used in this study: STEM21, STEM38, STEM86, and UCAL38. For the STEM lines, the collection and use of the human skeletal tissues was reviewed and approved by the Providence St. Joseph’s and St. Michael’s Healthcare Research Ethics Board (REB# 13-370) and the University of Toronto Office of Research Ethics reviewed the approved study and further assigned administrative approval (Protocol# 30754). Written consent was obtained from donors prior to the scheduled surgical procedure. Human skeletal muscle tissues removed from the multifidus muscle of patients undergoing lumbar spine surgery and designated for disposal were utilized in this study. Recruitment began August 14, 2014 and ended Aug 18, 2020. For the UCAL38 cell line, the collection and use of cadaveric human skeletal tissues was reviewed and approved by the University of Calgary Research Ethics Board (REB# 15-0005) and the University of Toronto Office of Research Ethics reviewed the approved study and further assigned administrative approval (Protocol# 37165). Written consent was from next of kin. Recruitment began Sept 14, 2018 and is ongoing. All procedures in this study were performed in accordance with the guidelines and regulations of the respective Research Ethics Boards. Table 1 summarizes muscle donors’ information for each line.

**Table 1.**
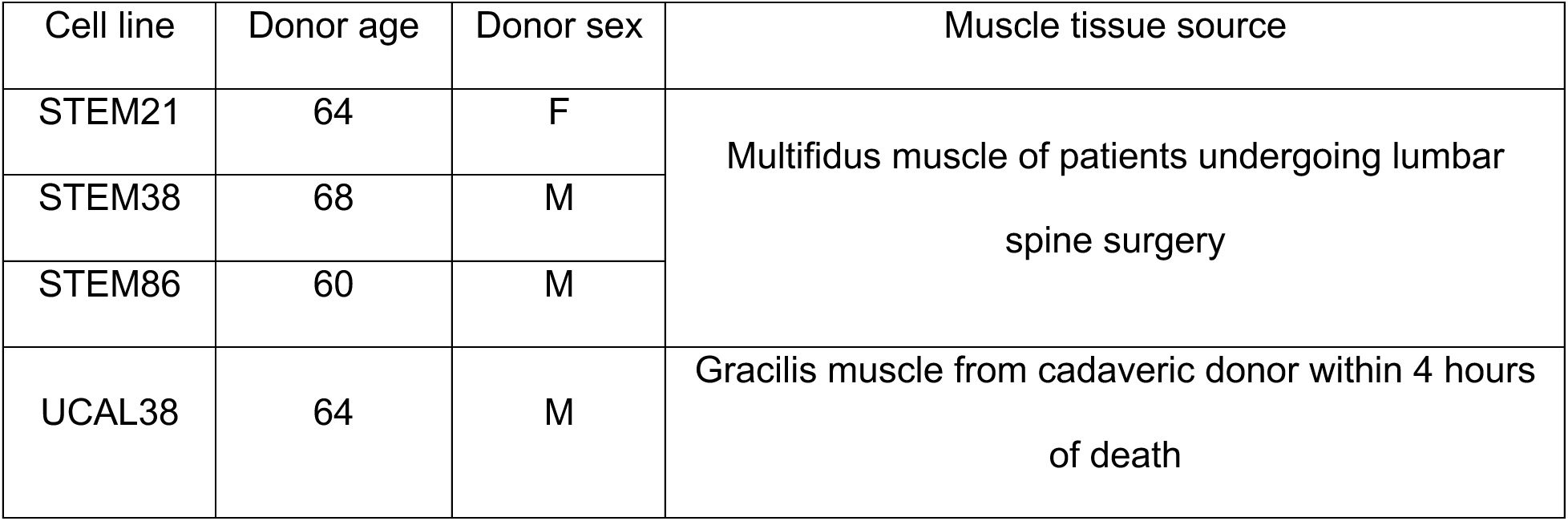
Skeletal muscle biopsy donor information.

| Cell line | Donor age | Donor sex | Muscle tissue source |
| --- | --- | --- | --- |
| STEM21 | 64 | F | Multifidus muscle of patients undergoing lumbar spine surgery |
| STEM38 | 68 | M |  |
| STEM86 | 60 | M |  |
| UCAL38 | 64 | M | Gracilis muscle from cadaveric donor within 4 hours of death |

The establishment of primary human myoblast cell lines was described in previous reports[30,31]. To summarize, human skeletal muscle biopsies were minced and incubated with collagenase (Sigma, 630 U/mL) and dispase (Roche, 0.03 U/mL) in Dulbecco’s Modified Eagle’s medium (DMEM; Gibco) to digest the ECM and basal lamina. The resulting cell slurry was then passed through a 20G needle multiple times to obtain a single-cell suspension, followed by incubation with red blood cell lysis buffer (Table 2). The resulting non-purified cells were plated on collagen-coated tissue culture dishes for one passage in myoblast growth medium (Table 2) and then were immunostained and sorted using fluorescence-activated cell sorting (FACS), to purify the CD56+ myogenic progenitor population.

**Table 2.** Cell culture media and solutions.

| Media and solutions | Composition |
| --- | --- |
| RBS lysis buffer | ddH<sub>2</sub>O, 15.5 mM NH<sub>4</sub>Cl (Sigma-Aldrich, A9434), 1 mM KHCO<sub>3</sub> (Sigma-Aldrich, 237205), 10 μM EDTA |
| Growth media | Ham's F-10 Nutrient Mixture (Wisent Bioproducts, 318-050-CL), 20% FBS (Gibco, 10437028), 5 ng/ml fibroblast growth factor-2 (ImmunoTools, 11343625), and 1% penicillin-streptomycin |

Myoblasts were expanded on collagen-coated tissue culture plastic plates before being collected at passage 7-8 for experiments using the following method: adherent cells were first washed with pre-warmed PBS, then incubated in 0.05 % trypsin-EDTA (Thermo Fisher Scientific, 25200072) at 37 °C until all cells detached (about 5 minutes). After neutralizing trypsin-EDTA with growth medium, the cell solution was centrifuged in 15 ml conical tubes at 300 G for 8 minutes, then resuspended in 1 ml fresh growth medium. After cell-counting using a hemocytometer, 20,000 cells were seeded per hydrogel (diameter=12mm) for morphology, focal adhesion analysis, and traction force microscopy experiments, while 25,000 cells were seeded per hydrogel for all other experiments.

### ML7 and LPA treatment

Prior to NE wrinkling analysis, cells were treated with 25 µM ML7 for 1 hour before fixing, or with 50 µM LPA for 2 hours post-seeding, followed by a wash with media before incubating cells in fresh growth media until the fixation time-point. Prior to Ki67 analysis, treatment with 10 µM ML7 or LPA was conducted overnight. In all of these conditions, the cells were fixed at 45 hours post-seeding.

### Immunostaining

Fixation was done by replacing half the volume of media in the culture dish with 4 % paraformaldehyde (Electron Microscopy Sciences; 15714) in phosphate buffered saline (PBS) followed by a 15-minute incubation. After three 5-minute PBS washes, cells were permeabilized in 0.1 % Triton-X100 (BioShop, TRX777.500)/PBS for 15 minutes. Cells were then blocked in 10 % goat serum in PBS for 1 hour. Immediately after blocking, primary antibodies diluted in 1 % goat serum were added and left overnight at 4 °C. The following day, after three 5-minute washes in 0.025 % Tween-20 (BioShop, TWN510.500)/PBS, cells were incubated in secondary antibodies and Hoechst counterstain diluted in 1 % goat serum for 1 hour. For F-actin visualization, Phalloidin was also added at this step. Antibodies information and dilution are reported in Table 3. Before imaging, cells were washed for 5 minutes in 0.0025 % Tween-20/PBS, then a final wash on a gentle shaker for 30 minutes. All steps were done at room temperature unless specified otherwise. For Pax7/MyoD experiments, after the final wash, PBS was removed then a coverslip was placed on top of the cells, so that imaging was done through the top coverslip with the plate upside-down in case PAA hydrogels with different stiffness alter the perceived fluorescence intensity of the samples, even though this has not been reported as a concern in major PAA protocols[29,32]. In other experiments where fluorescence intensity was not critical, imaging was done through the bottom coverslip of the glass-bottom plate.

**Table 3.** Antibody information.

| Antibody | Host species | Dilution ratio | Manufacturer, catalog number |
| --- | --- | --- | --- |
| Hoechst 33342 | - | 1:1000 | Thermo Fisher Scientific, H3570 |
| Fluorescein Phalloidin | - | 1:400 | Molecular Probes, F432 |
| anti-Paxillin, clone 5H11 | mouse IgG1 | 1:200 | Millipore Sigma, 05-417 |
| anti-Pax7 | mouse<br>IgG1 | 1:1 | In-house supernatant from<br>hybridoma cell line (DHSB) |
| anti-MyoD1 | mouse<br>IgG2b | 1:300 | Santa Cruz Biotechnology, sc-377460 |
| anti-Ki67 | rabbit | 1:300 | Abcam, ab16667 |
| anti-Lamin A/C | mouse<br>IgG1 | 1:300 | Santa Cruz Biotechnology, sc-376248 |
| anti-Lamin A/C (H-110) | rabbit | 1:300 | Santa Cruz Biotechnology, sc-20681 |
| Alexafluor™ 488 Anti-mouse<br>IgG (H+L) | goat | 1:300 | Invitrogen, A11001 |
| Alexafluor™ 647 Anti-mouse<br>IgG1 | goat | 1:300 | Invitrogen, A21240 |
| Alexafluor™ 546 Anti-mouse<br>IgG (H+L) | goat | 1:300 | Invitrogen, A11003 |
| Alexafluor™ 546 Anti-mouse<br>IgG2b | goat | 1:300 | Invitrogen, A21141 |
| Alexa Fluor® 647 Mouse Anti-Human CD56 (NCAM-1) | mouse<br>IgG1 | 1:20 | BD Pharmingen, 557711 |

**Table 4.**
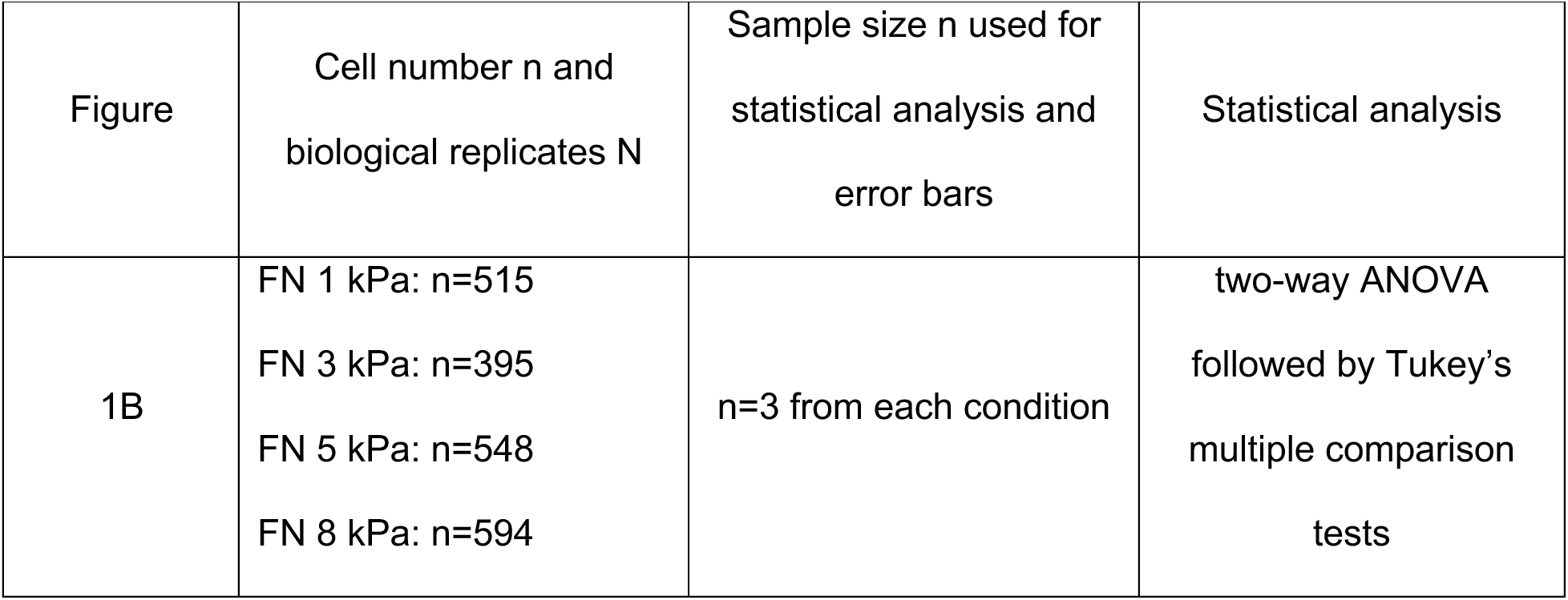
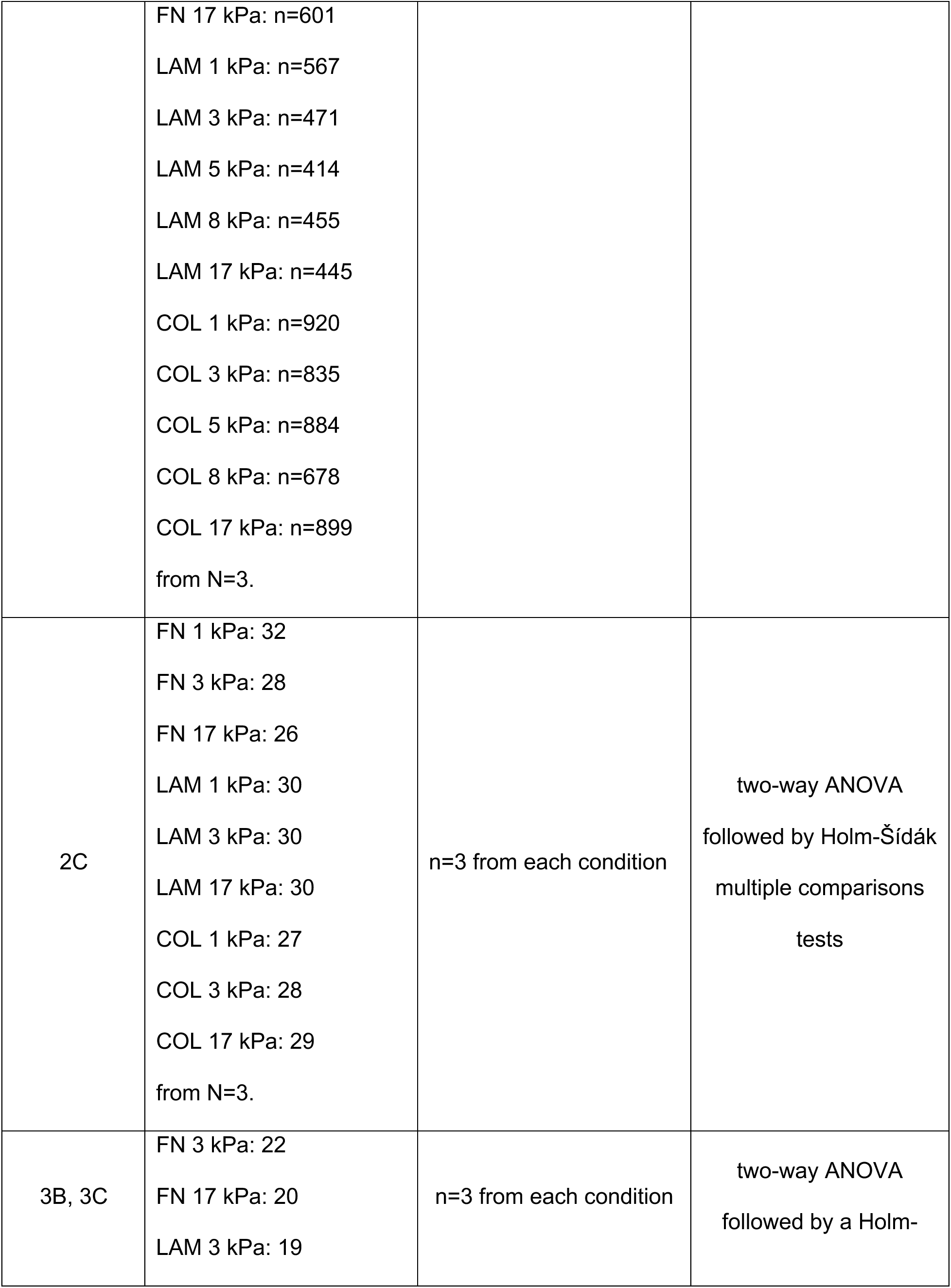
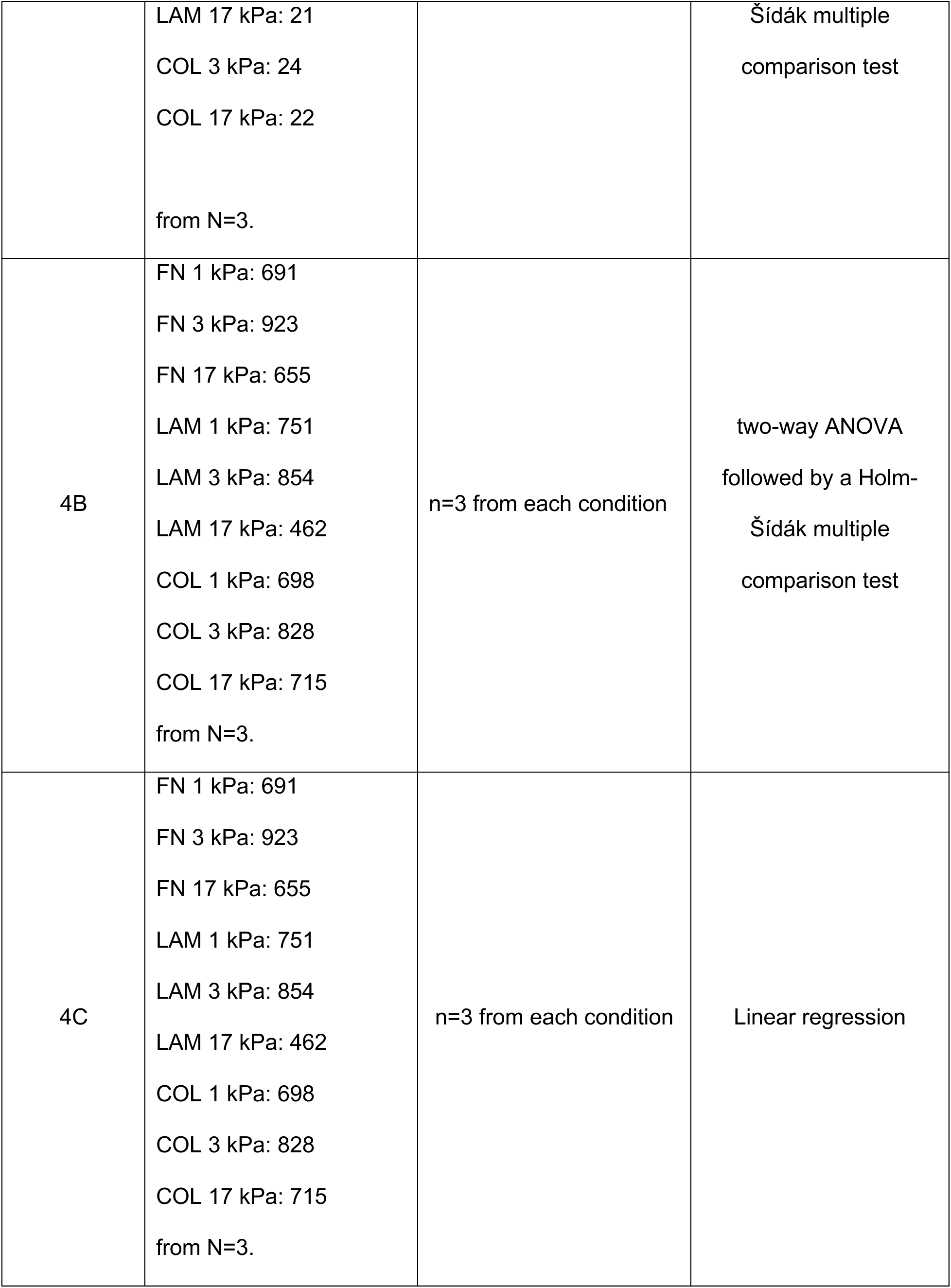
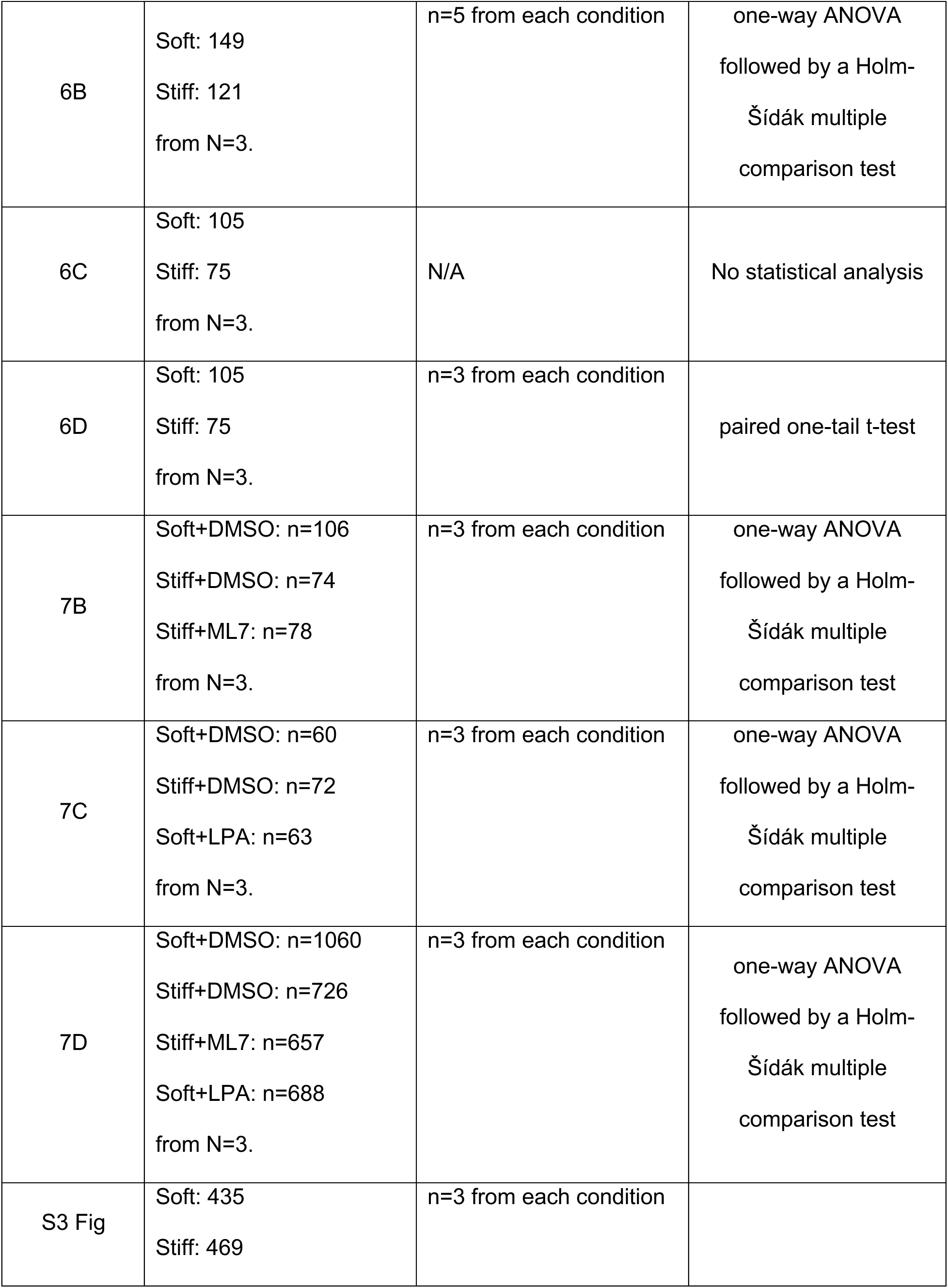
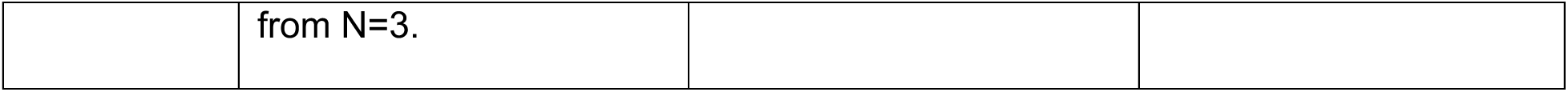
Experimental replicate breakdown and statistical analysis.

| Figure | Cell number n and biological replicates N | Sample size n used for statistical analysis and error bars | Statistical analysis |
| --- | --- | --- | --- |
| 1B | FN 1 kPa: n=515<br>FN 3 kPa: n=395<br>FN 5 kPa: n=548<br>FN 8 kPa: n=594 | n=3 from each condition | two-way ANOVA followed by Tukey's multiple comparison tests |
|  | FN 17 kPa: n=601<br>LAM 1 kPa: n=567<br>LAM 3 kPa: n=471<br>LAM 5 kPa: n=414<br>LAM 8 kPa: n=455<br>LAM 17 kPa: n=445<br>COL 1 kPa: n=920<br>COL 3 kPa: n=835<br>COL 5 kPa: n=884<br>COL 8 kPa: n=678<br>COL 17 kPa: n=899<br>from N=3. |  |  |
| 2C | FN 1 kPa: 32<br>FN 3 kPa: 28<br>FN 17 kPa: 26<br>LAM 1 kPa: 30<br>LAM 3 kPa: 30<br>LAM 17 kPa: 30<br>COL 1 kPa: 27<br>COL 3 kPa: 28<br>COL 17 kPa: 29<br>from N=3. | n=3 from each condition | two-way ANOVA<br>followed by Holm-Šídák<br>multiple comparisons<br>tests |
| 3B, 3C | FN 3 kPa: 22<br>FN 17 kPa: 20<br>LAM 3 kPa: 19 | n=3 from each condition | two-way ANOVA<br>followed by a Holm- |
|  | LAM 17 kPa: 21<br>COL 3 kPa: 24<br>COL 17 kPa: 22<br>from N=3. |  | Šídák multiple comparison test |
| 4B | FN 1 kPa: 691<br>FN 3 kPa: 923<br>FN 17 kPa: 655<br>LAM 1 kPa: 751<br>LAM 3 kPa: 854<br>LAM 17 kPa: 462<br>COL 1 kPa: 698<br>COL 3 kPa: 828<br>COL 17 kPa: 715<br>from N=3. | n=3 from each condition | two-way ANOVA<br>followed by a Holm-Šídák multiple comparison test |
| 4C | FN 1 kPa: 691<br>FN 3 kPa: 923<br>FN 17 kPa: 655<br>LAM 1 kPa: 751<br>LAM 3 kPa: 854<br>LAM 17 kPa: 462<br>COL 1 kPa: 698<br>COL 3 kPa: 828<br>COL 17 kPa: 715<br>from N=3. | n=3 from each condition | Linear regression |
| 6B | Soft: 149<br>Stiff: 121<br>from N=3. | n=5 from each condition | one-way ANOVA<br>followed by a Holm-Šídák multiple<br>comparison test |
| 6C | Soft: 105<br>Stiff: 75<br>from N=3. | N/A | No statistical analysis |
| 6D | Soft: 105<br>Stiff: 75<br>from N=3. | n=3 from each condition | paired one-tail t-test |
| 7B | Soft+DMSO: n=106<br>Stiff+DMSO: n=74<br>Stiff+ML7: n=78<br>from N=3. | n=3 from each condition | one-way ANOVA<br>followed by a Holm-Šídák multiple<br>comparison test |
| 7C | Soft+DMSO: n=60<br>Stiff+DMSO: n=72<br>Soft+LPA: n=63<br>from N=3. | n=3 from each condition | one-way ANOVA<br>followed by a Holm-Šídák multiple<br>comparison test |
| 7D | Soft+DMSO: n=1060<br>Stiff+DMSO: n=726<br>Stiff+ML7: n=657<br>Soft+LPA: n=688<br>from N=3. | n=3 from each condition | one-way ANOVA<br>followed by a Holm-Šídák multiple<br>comparison test |
| S3 Fig | Soft: 435<br>Stiff: 469 | n=3 from each condition |  |
|  | from N=3. |  |  |

### Traction force analysis

After 16 hours of culture on PAA hydrogels with embedded fluorescent beads (ThermoFisher Scientific, F8800), the cells were stained for CellTracker Deep Red dye (ThermoFisher Scientific, C34565) according to manufacturer’s protocol. Imaging was done in FluoroBrite DMEM (ThermoFisher Scientific, A1896701), 20 % FBS (Gibco, 10437028), and 1 % penicillin-streptomycin solution medium. Images of beads and cells were taken before and after the addition of 10 % sodium dodecyl sulphate (SDS). Traction stress maps were obtained using ImageJ macros developed in prior published work [33,34]. Using MATLAB, cell masks were obtained by expanding the original CellTracker binarized image by 70 pixels in all directions. Only traction stress values inside the cell masks were considered.

### EdU assay

After 16 hours of culture on hydrogel substrates, cells were pulsed with EdU at a concentration of 1 µM for 12 hours. Cells were fixed in 4 % PFA for 15 minutes and incubated in blocking solution containing 10 % goat serum and 0.3 % Triton X-100 (BioShop, TRX777) diluted in PBS for 45 minutes at room temperature. EdU immunolabeling was done using the Invitrogen Click-iT EdU kit according to the manufacturer’s protocol (Invitrogen™, C10339) and then cells were counterstained with Hoechst 33342.

### Viability assay

After 16 hours of culture on hydrogel substrates, cells were washed with PBS, then incubated with 1 µM calcein-AM (ThermoFisher Scientific, L3224), 1 µM propidium iodide (Sigma, P4864) or ethidium homodimer-1 (ThermoFisher Scientific, L3224), and 1:1000 Hoechst in PBS for 20 minutes at room temperature. After removing the staining solution, cells were washed with PBS and imaged immediately.

### Microscopy and image analysis

Except for the EdU analysis, all microscopy was done with a Zeiss Axio Observer 7 microscope equipped with LSM 800 scan head. A Plan-APO 40x/NA 1.40 Oil DIC objective was used for all single cell analyses (i.e. paxillin, lamin A/C, traction force microscopy, and phalloidin representative images), and a Plan-Apo 10x/NA 0.45 objective was used for cell spreading area and the Ki67 imaging. EdU staining images were taken using an Olympus IX83 microscope equipped with a DP80 dual CCD camera and either a LUCPLFLN PH 20x/ NA 0.46 or UPLFLN 2PH 10x/ NA 0.3 objective.

All image analysis was performed with ImageJ. FA alignment by goodness of fit (GoF) to a Gaussian curve was obtained by running the Directionality ImageJ plug-in (written by Jean-Yves Tinevez; https://imagej.net/plugins/directionality) on a random region of interest (ROI) close to the cell periphery on the maximum projection of paxillin z-stacks. The analysis performed was based on Fourier spectrum analysis. The percentage of total cells labeled by EdU was quantified by dividing the number of segmented objects in the EdU channel by the number of segmented objects in Hoechst channel. Nuclear wrinkling index was quantified using the ImageJ macro developed by Cosgrove et. al [35] with some adjustments to allow semi-batch processing of our datasets. The adapted macro used can be found in our public Github repository (https://github.com/gilbertlabcode/Nguyen_PLOS_One2023). Ki67, Pax7, and MyoD fluorescence intensities in each cell were quantified by measuring the integrated density inside each nucleus. Binary binning (+/-) of Ki67, Pax7, and MyoD was done by identifying an intensity threshold below which cells appear “negative” by reference to the secondary antibody controls.

### Lentiviral production

Lentivirus particles were produced by calcium phosphate transfection, as reported previously[36]. Briefly, 35,000/cm^2^ HEK 293T cells were seeded in a 10-cm culture plate and co-transfected 2 days later at 80 % confluency with second generation lentivirus system. Culture media (DMEM supplemented with 10 % FBS) was refreshed prior to transfection, followed by addition of the plasmid cocktail containing 4 µg of pMD2.G (a gift from Didier Trono; Addgene plasmid # 12259; http://n2t.net/addgene:12259; RRID:Addgene_12259), 10 µg of psPAX2 (a gift from Didier Trono; Addgene plasmid # 12260; http://n2t.net/addgene:12260; RRID:Addgene_12260) and 15 µg of pLVX-EF1a-GFP-LaminA-IRES-Hygromycin [37] (a gift from David Andrews; Addgene plasmid # 134867; http://n2t.net/addgene:134867; RRID:Addgene_134867) for 12 hours. Culture media was replaced to OptiMEM (Thermofisher, 31985062) 20mM HEPES (Sigma-Aldrich, 51558-50ML), and the supernatant containing viral particles was collected at 24 and 48 hours after transfection, passed through a 0.45 µm filter, and then concentrated using a 50 mL 100 000 MWCO spin column (Cytiva Life Sciences).

### Lentiviral transduction

Primary human myoblasts were plated in a collagen-coated 6-well plate in growth media and transduced when the cells were at about 50% confluency (i.e. ∼ 24 hours after plating). 150-500 µL of viral supernatant containing 6 µg/mL polybrene (EMD Millipore, TR-1003-G) and was added dropwise to the cells in fresh growth media and then incubated for 12 hours. The virus containing media was aspirated and the cells were returned to regular growth media for 48 hours. The cells were then trypsinized and seeded onto 17 kPa hydrogel substrates for 16 hours until traction force microscopy experiment.

### Proteomics

#### Reagents

Trypsin from bovine pancreas (Promega Biotec, V5113), lysine-C (New England BioLabs, P8109S), protease and phosphatase inhibitor cocktail (ThermoFisher Scientific, 78442), formic acid (FA, Sigma–Aldrich, F0507-100ML), tris(2-carboxyethyl) phosphine (TCEP, Sigma–Aldrich, C4706-2G), iodoacetamide (IAA, Sigma–Aldrich, I1149-5G), acetonitrile (≥ 99.9 %, CAN, Fisher Chemicals, A9551), and water (Fisher Chemicals, W6-4).

#### Preparation and digestion of protein extracts from myoblasts

Myoblasts cultured on hydrogel substrates for 45 hours were collected by incubation with trypLE™ (ThermoFisher Scientific, 12604013). Cell numbers were determined by hemocytometer counting. For each stiffness condition, a total number of 100,000 cells were washed three times with cold PBS and resuspended in 20 μl lysis buffer (50 mM PBS, 1 % (v/v) protease inhibitor cocktail). After three continuous freezing and defrost cycles, insoluble portions were separated from the soluble portions by centrifugation at 16,000 × g for 20 minutes at 4 °C. Samples were reduced in 10 mM TCEP (final concentration) at 70 °C for 30 minutes, alkylated with 20 mM IAA (final concentration) in the dark at room temperature for 30 minutes, followed by incubation with lysine-C (1:60) at 37 °C for 1.5 hr. Trypsin (1: 25) was then added and incubated at 37 °C for 16 hr. Finally, digests were quenched by the addition of 1 μL of concentrated formic acid.

#### NanoRPLC-ESI-MS/MS methods

Nanoflow reversed-phase liquid chromatography (NanoRPLC) was performed on an EASY-nLC 1200 ultra-high-pressure system coupled to a Q-Exactive HF-X mass spectrometer equipped with a nano-electrospray ion source (Thermo Fisher Scientific). The obtained peptides were automatically loaded onto a C18 trap column (100 μm i.d. × 5 cm) and separated by a C18 capillary column (100 μm i.d. × 15 cm). The trap column and the analytical column were both packed in-house with 1.9 μm, 120 Å ReproSil-Pur C18 reversed phase particles (Dr. Maisch GmbH). Two mobile phases (A: 0.1 % (v/v) FA and B: 80/20/0.1% ACN/water/formic acid (v/v/v) were used to generate a 130 minutes gradient with the flow rate of 300 nL/minute (kept at 3 % B for 5 minutes, 90 minutes from 3 % to 30 % B, 20 minutes from 30 % to 45 % B, 1 minutes from 45 % to 95 % B and 14 minutes to kept at 95 % B). MS was operated in full scan (120,000 FWHM, 400-1100 m/z) and data-independent acquisition (DIA) modes at positive ion mode. A 25.0 m/z isolation window was set for MS/MS scan (60,000 FWHM) with stepped HCD collision energy of 25.5 %, 27 % and 30 %. The maximum IT for both MS and MS/MS was 254 ms. The electro-spray voltage was 2.0 kV, and the heated capillary temperature was 275 °C. All the mass spectra were recorded with Xcalibur software (version 4.1, Thermo Fisher Scientific).

#### Database search

DIA-NN 1.8 were applied for database searching against the Uniprot_Homo Sapiens database (updated on 11/11/2021, 78120 entries) and the predicted spectrum library. The predicted spectrum library was generated by DIA-NN 1.8 using the same database. The parameters of database searching include up to one missed cleavage allowed for full tryptic digestion, precursor ion mass tolerance and product ion mass tolerance were set 10 and 20 ppm, respectively, carbamidomethylation (C) as a fixed modification, and oxidation(M), acetyl (protein N-term) as variable modifications. Match between runs (MBR) was activated. Peptide spectral matches (PSM) were validated based on q-value at a 1 % false discovery rate (FDR).

#### Data processing

Overrepresentation tests were done with PANTHER database version 17.0 released on 2022-02-22 using Fisher’s exact test with false discovery rate correction. For PANTHER pathways annotation, the input was a list of differentially regulated proteins in both directions. For GO cellular component annotation, the inputs were upregulated proteins in either soft or stiff conditions.

### Data normalization

Cell spreading area: Cell area data had a log-normal distribution (S1 A-B Fig). To meet the normality assumption underlying the ANOVA test to compare the means between substrate conditions, analysis was done based on the natural log of cell area (ln(Area)). Additionally, our experiments were repeated on primary human myoblasts lines from independent donors. To minimize cell line variance that may mask substrate effects, we normalized the mean ln(Area) values to that of the values obtained on the 9 kPa condition for each cell line: normalized ln(Area) = ln(Area)/ln(Area)_8kPa_. Pre-normalized data is reported in S1 C-E Fig.

EdU assay: aside from reporting the unadjusted data, we scaled each cell line by the maximum % total EdU+ value (scaled x = x/x_max_) before linear regression fitting. Normalization by scaling allowed us to focus on comparing the stiffness-dependent trends and minimized cell line variance in the exact values of % total EdU+ cells.

### Statistical analysis

All statistical analysis was conducted using GraphPad Prism 6.0 software. To compare two conditions in experiments with one independent variable, comparisons were done with t-tests. To compare more than two conditions in experiments with one independent variable, a one-way ANOVA test preceded Holm-Šídák multiple comparison tests. For experiments with two independent variables, two-way ANOVA preceded Holm-Šídák multiple comparison tests. For all tests, α=0.05. p-values used to discuss comparisons between conditions are reported on top of graphs.

### Representative image preparation

Figure 1: Cropped images were placed on a black background separated by white lines. Images were max projection of z-stacks.

Figure 2: Raw paxillin staining images were processed according to a published process [38] up to the Log3D filtering step to create representative images. These images were then processed with the Log3D plug-in, binarized, and cropped to create the insets.

Figure 3: Filtered stress maps were created following the process written in the “Traction force analysis” section above.

Figure 4: The brightness and contrast of raw EdU and Hoechst images were equally adjusted in all conditions.

Figure 5: Individual nuclei were cropped from raw Lamin A/C images. The brightness and contrast of Lamin A/C were equally adjusted for all nuclei.

Figure 6: Images from the first three conditions (soft+DMSO, stiff+DMSO, stiff+ML7) were collected from the same experiment so their brightness and contrast were adjusted equally. The images for the soft+LPA condition were collected from a separate experiment, and thus, the brightness and contrast were adjusted to most closely match that of the other conditions. Note that fluorescence intensity was not considered in the analysis of these results.

## Results

### Accentuated myoblast stiffness sensitivity observed on fibronectin-tethered substrates

We first explored the influence of ECM ligand tethering on human primary myoblast responses associated with stiffness-sensing that included cell spreading area, focal adhesion assembly, and traction stress. We followed a well-established poly-acrylamide (PA) hydrogel fabrication protocol to create cell culture substrates with tuned stiffness that we then tethered with ECM ligands found within the native muscle stem and progenitor cell microenvironment: fibronectin (FN), laminin (LAM), or collagen-1 (COL)[29]. We selected PA gel stiffnesses of 1, 3, 5, 8, and 17 kPa to mimic the reported range of the Young’s modulus of skeletal muscle and single muscle fibers, which were reported to be approximately 0.2-18 kPa, spanning healthy, damaged, and aged states [4,13,20,21,39,40].

16 hours after plating on FN substrates, human primary myoblast cell spreading was minimal on 1 kPa culture substrates, noticeably increased at 3kPa, then plateaued from 3 to 17 kPa (Fig 1). Cell spreading on LAM substrates presented a similar relationship with culture substrate stiffness, although a slight decrease in spreading at 17 kPa was observed. By contrast, cells on COL substrates attained equally large spreading area independent of culture substrate stiffness within the tested stiffness range, except for a decrease at 17 kPa (Fig 1A-B). As we only included well isolated mononucleated cells in our analysis, the decrease in spreading area at 17 kPa may have been an analysis artifact due to the greater likelihood of larger cells making physical contact with neighboring cells.

**Fig 1.**
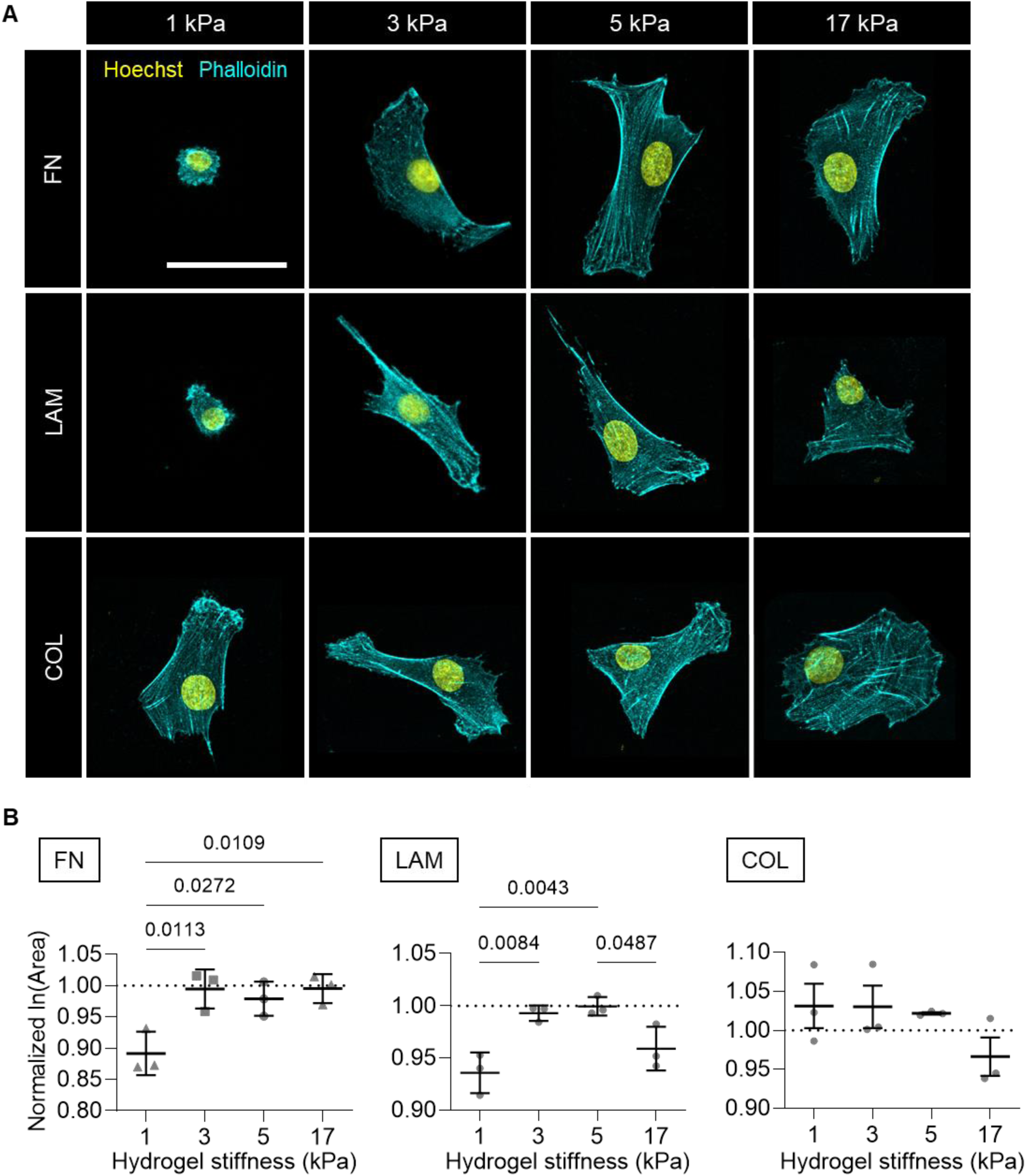
Cell spreading area characterization across substrate stiffness and ECM conditions. **A** Representative confocal images of phalloidin staining in primary human myoblasts cultured for 16 hours on 1, 3, 5, or 17 kPa polyacrylamide gels tethered with fibronectin (FN), laminin (LAM), or collagen 1 (COL). Phalloidin-cyan, Hoechst-yellow. scale bar=50 µm **B** Graphs showing mean cell spreading area across hydrogel stiffness, in FN, LAM, and COL conditions, in terms of fold change with respect to the 8 kPa condition (dotted line at y=1). Each data point represents the mean of one biological replicate. n=395-920 cells per condition across N=3 biological replicates. Error bars report mean ± SD. Statistical comparisons were made by two-way ANOVA followed by Tukey’s multiple comparison tests.

Next, we visualized adhesion structures with paxillin immunostaining and noted that the paxillin structures detected had a broad range of sizes (Fig 2A). On the 1 kPa substrates tethered with FN and LAM, wherein the myoblasts had minimal spreading, we observed long paxillin+ structures outlining the nucleus and/or the peripheral edges of the cells. The appearance of cells with these features was rare on the higher substrate stiffnesses and were also rare on the COL tethered substrates more generally. Adhesion structure morphologies varied along three scales: within a single cell, across different cells cultured on the same substrates, and between the biological replicate cell lines. These variabilities made it challenging to attribute substrate effects to adhesion size and shape, which, by themselves, are insufficient to infer focal adhesion function and maturity [41]. For this reason, we instead chose to quantify local adhesion structure alignment. Since cells exert traction forces through adhesion structures where stress fibers terminate, the alignment of adhesion structures is indicative of the alignment of stress fibers and provides a speculative imagery of the cell contractility state[42,43]. To measure the alignment of adhesion structures, we used the “Directionality” ImageJ plugin, which generates a frequency distribution histogram of structures over a range of direction angles (with 0° being the East direction and going counterclockwise). The histogram was then fitted to a Gaussian curve. If the structures in the region of interest align in a common direction, the histogram will have a peak and resemble a Gaussian curve, thus having a high goodness of fit (GoF) (Fig 2B, top). Conversely, if a common direction of alignment does not exist, the histogram will be flat and have a low GoF (Fig 2B, bottom). The alignment of paxillin structures across substrate conditions showed trends like the cell spreading area data (Fig 1). Specifically, on FN and LAM substrates, GoF was low at 1kPa, increased at 3kPa and then remained relatively constant at 17kPa, although the increase on LAM substrates was more subtle, and did not reach statistical significance in our study (Fig 2C). On the COL substrates, adhesion structures reached equally high alignment regardless of substrate stiffness (Fig 2C).

**Fig 2.**
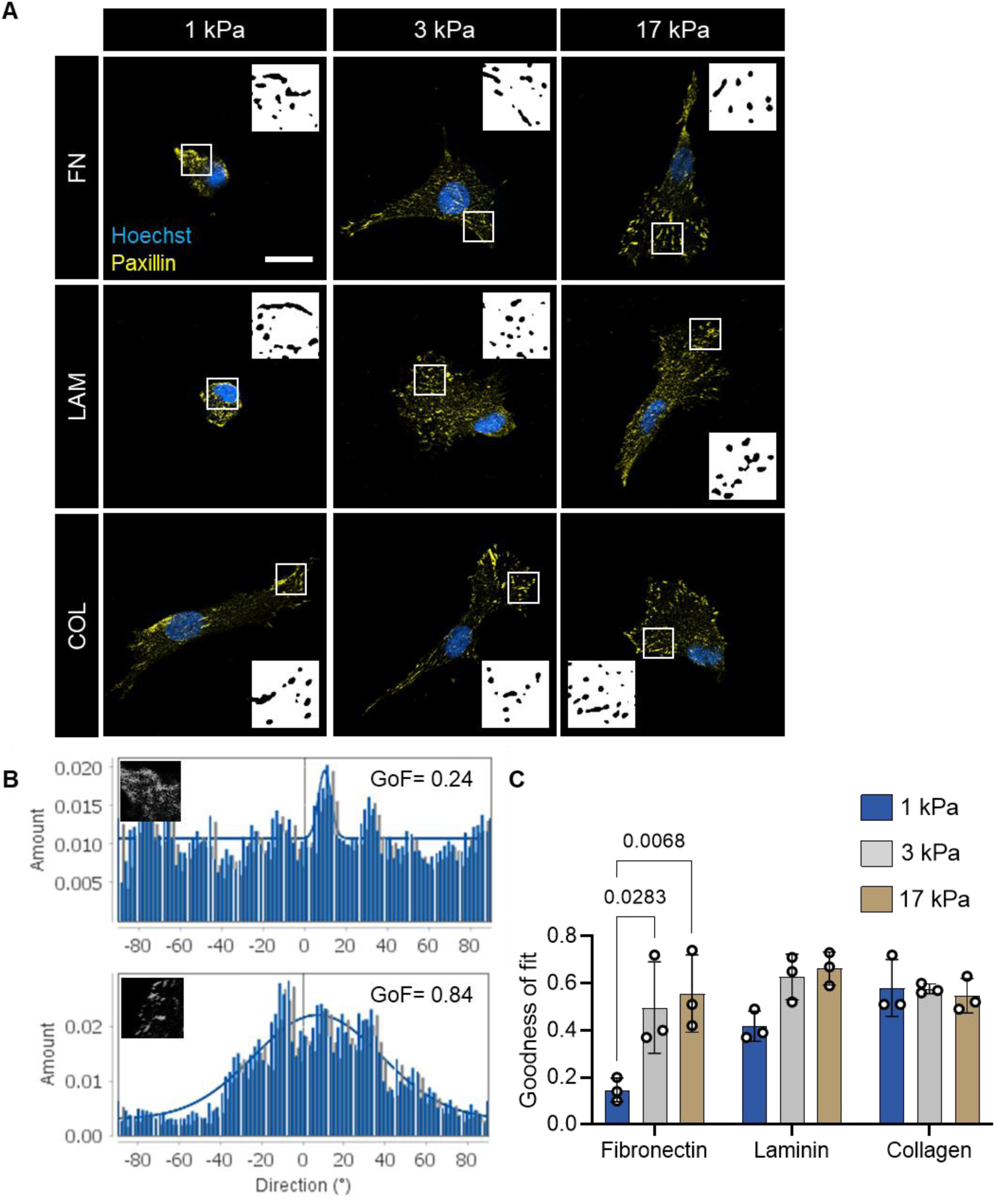
Adhesion alignment characterization across substrate stiffness and ECM conditions. **A** Representative confocal images of primary human myoblasts adhesion structures 16 hours post-seeding on fibronectin (FN), laminin (LAM) or collagen 1 (COL) tethered polyacrylamide gels. Paxillin-yellow, Hoechst-blue. Scale bar=20 μm. **B** Representative directionality histograms of a paxillin staining ROI with goodness of fit to a Gaussian curve (GoF) = 0.24 (top) and an ROI with GoF=0.84 (bottom). y-axis is the frequency of structures detected by the algorithm and x-axis is the angle direction of those structures with respect to a horizontal line (°) **C** Bar graph showing GoF as a metric of adhesion alignment. Each data point represents the mean of one biological replicate. n=26-32 cells per condition across N=3 biological replicates. Error bars report mean ± SD. Statistical comparisons were made by two-way ANOVA followed by Holm-Šídák multiple comparisons tests.

Although cell spreading area and adhesion alignment did not differ significantly between 3 and 17 kPa on FN substrates, upon conducting traction force microscopy studies, we observed a significant increase in mean and max traction stress across these conditions. Similar trends were seen on the LAM and COL substrates, though the magnitude of the effect was more subtle than on the FN condition (Fig 3A-C).Traction data on 1 kPa substrates were excluded from this analysis because we were unable to reliably derive cellular traction stress with our workflow because of the extremely large hydrogel deformations. Likely due to large deformation caused on softer hydrogels, even though lower stresses were recorded on 3 kPa substrates compared to 17 kPa, strain energy was generally higher on 3 kPa than 17 kPa in all ECM conditions, but was only statistically significant for COL (S3 Fig).

**Fig 3.**
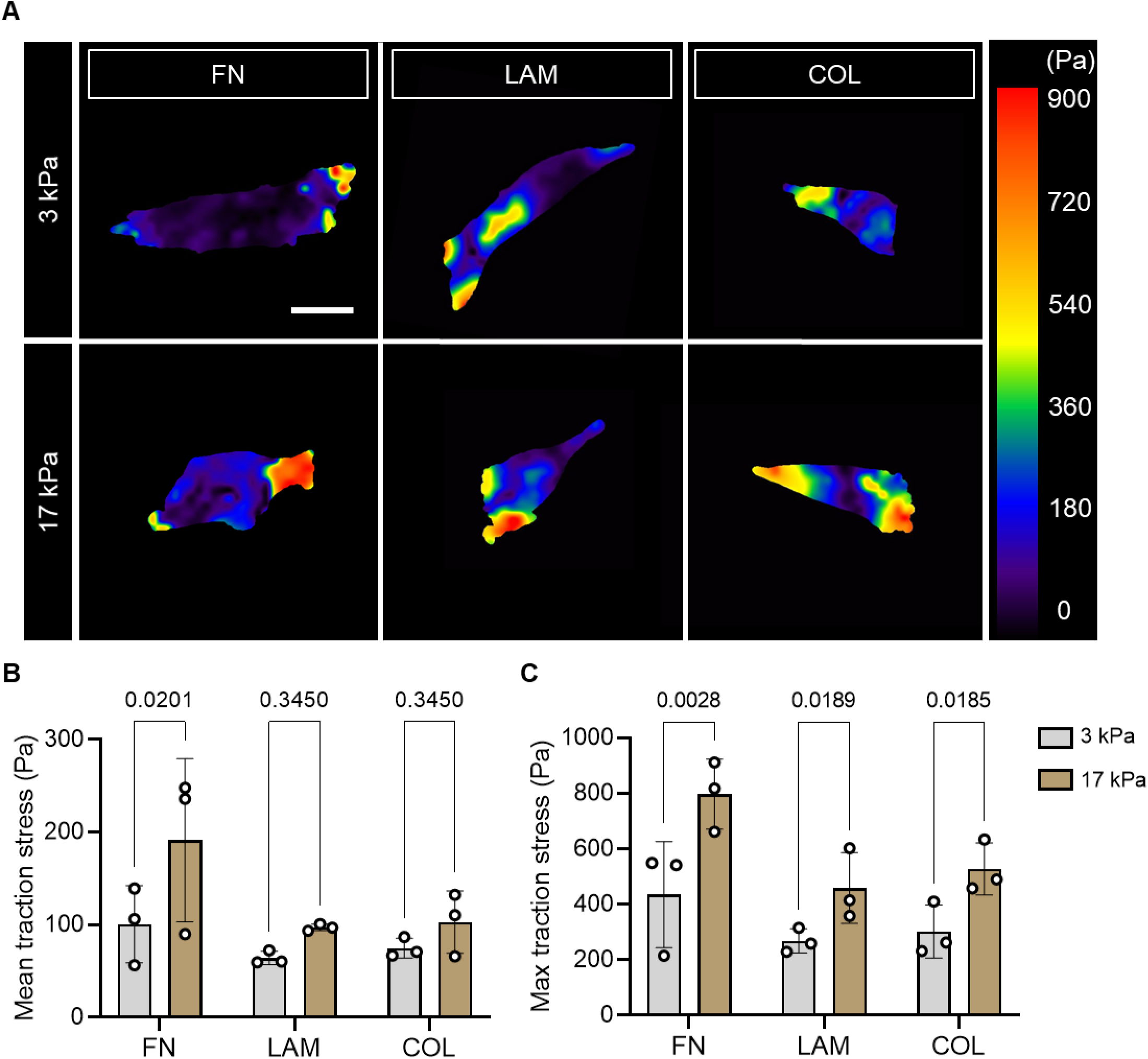
Cellular traction stress measurement across substrate stiffness and ECM conditions. **A** Representative traction stress maps of myoblasts 16 hours post-seeding on fibronectin (FN), laminin (LAM) or collagen 1 (COL) tethered polyacrylamide gels. Scale bar=20 μm **B and C** Bar graphs showing mean and max traction stress, respectively, across substrate stiffness and ECM conditions. n=19-24 cells per condition across N=3 biological replicates. Each data point represents the mean of one biological replicate. Error bars report mean ± SD. Statistical comparisons were made by two-way ANOVA followed by a Holm-Šídák multiple comparison test.

In summary, by evaluating the morphology, focal adhesion arrangement, and cellular traction stress of primary human myoblasts cultured on 1, 3, and 17kPa PA hydrogels, we found that FN-tethered hydrogel substrates encouraged more pronounced stiffness sensitivity when compared against the LAM- and COL-tethered substrates, in terms of the above metrics.

### ECM ligands influence the linearity of proliferation correlation with substrate stiffness

To probe whether ECM ligands also tune cellular functional response to substrate stiffness, we evaluated myoblast proliferation using the 5-ethynyl-2’-deoxyuridine (EdU) assay. After cells were cultured for 16 hours on hydrogel substrates, we added EdU in the culture media for 12 hours. The percentage of EdU+ cells increased steadily from 1 to 17 kPa for myoblasts cultured on FN-tethered substrates (Fig 4B). For LAM- and COL-tethered substrates, the percentage of EdU+ cells showed an insignificant, trended increase from 1 to 3 kPa and then remained constant from 3 to 17 kPa. We suspected that myoblast proliferation and substrate stiffness had a positive, linear correlation on FN but not on LAM and COL substrates. To assess linearity, we scaled %EdU+ values to minimize cell line variance while maintaining the correlation slope (refer to Methods) then performed linear regression fitting. On FN substrates, %EdU+ cells and substrate stiffness were indeed linearly fitted with an r^2^=0.7 while LAM and COL were poorly linear with r^2^ values of 0.3 and 0.2 respectively. The correlation slope in FN condition was also significantly non-zero with p=0.007 while p-values for LAM and COL were higher at 0.160 and 0.291 (Fig 4C). To eliminate the possibility that non-proliferative cells were instead undergoing apoptosis, we performed a live/dead assay, whose result indicated that all cells across all stiffnesses on FN-tethered substrates were calcein-AM positive and propidium iodide negative (S2 Fig).

**Fig 4.**
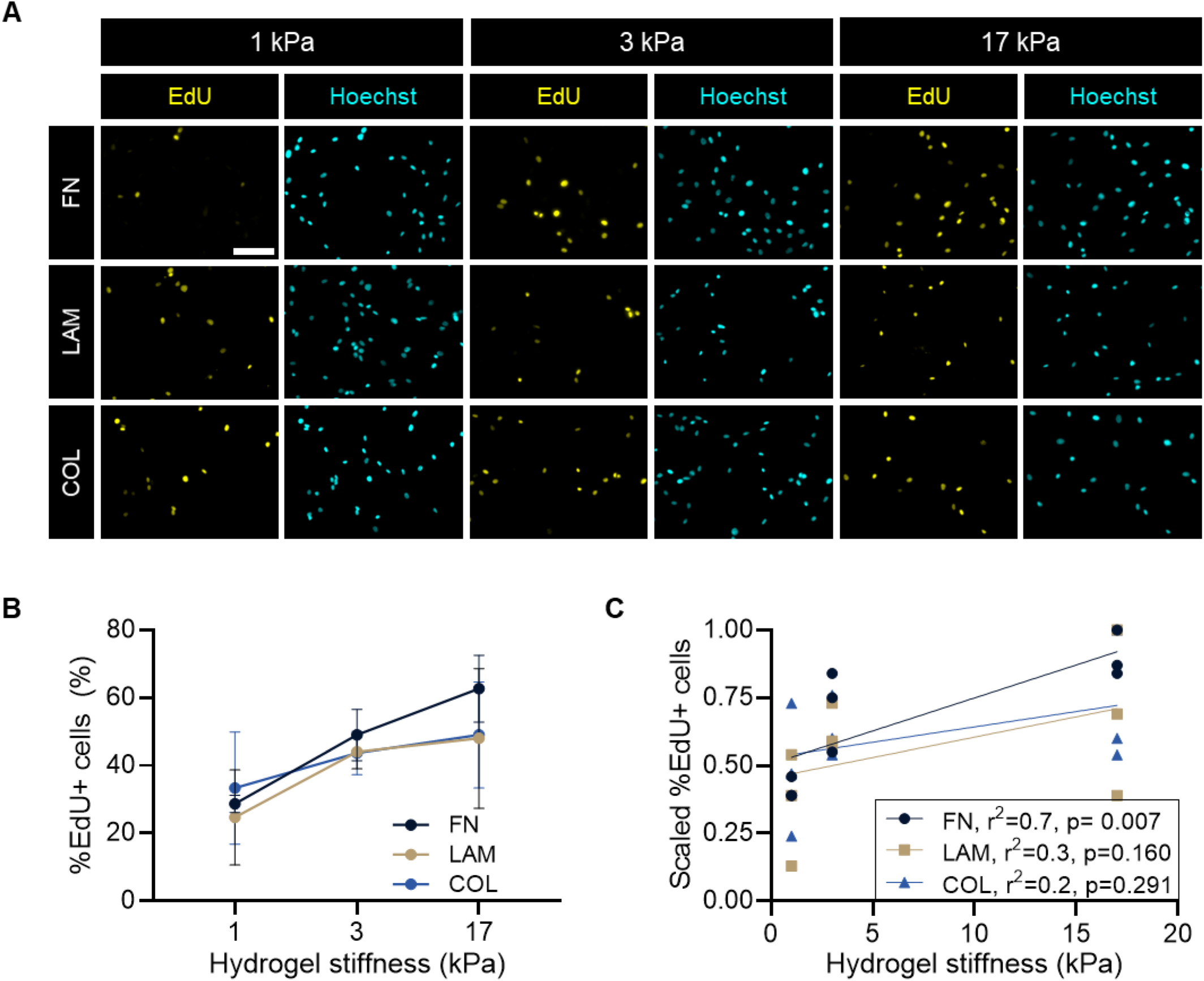
Myoblast proliferation exhibits a more linear correlation with stiffness on fibronectin (FN) substrates. **A** Representative immunostaining images of EdU incorporation assay at 16hr post-seeding on fibronectin (FN), laminin (LAM) or collagen 1 (COL) tethered polyacrylamide gels followed by a 12hr EdU pulse. EdU-yellow, Hoechst-cyan. Scale bar=100 μm. **B** Line graph showing the mean ± SD of % total EdU+ cells. Statistical comparisons were made by two-way ANOVA followed by Holm-Šídák multiple comparison test **C** Linear regression lines of the same data set in B. EdU percentages scaling was conducted by normalizing to the max value of each cell line in order to eliminate cell line variation (scaled x = x/x_max_). Legend displays r^2^ (goodness of fit) and p-value testing to determine whether the slope of the fitted line is significantly non-zero. Each data point represents the mean of one biological replicate. n=462-923 cells per condition across N=3 biological replicates.

### Nuclear envelope (NE) wrinkling can predict Ki67 expression

In conducting EdU analysis, we noticed qualitatively that there were more cell nuclei with odd shapes on softer substrates, so we asked if NE morphology regulation could be upstream of stiffness-dependent proliferation control. We followed through with a series of experiments focusing on two conditions: soft (1kPa) and stiff (17kPa) FN substrates. Immunostaining for Lamin A/C revealed a heterogeneity in NE shapes on both stiffnesses: some nuclei were taut and oval-like, as generally expected for adherent cells, but some were severely wrinkled (Fig 5A). We quantified NE wrinkling by measuring wrinkle index (WI) using an ImageJ macro developed by Cosgrove et al. with minor modifications[35]. We found that taut nuclei with no or minor folding have WI ≤ 20%, whereas wrinkled nuclei have WI > 20%. Noticeably, the percentages of wrinkled nuclei, defined by WI > 20%, were two times higher on soft compared to stiff FN-tethered substrates (around 40% and 19%, respectively).

**Fig 5.**
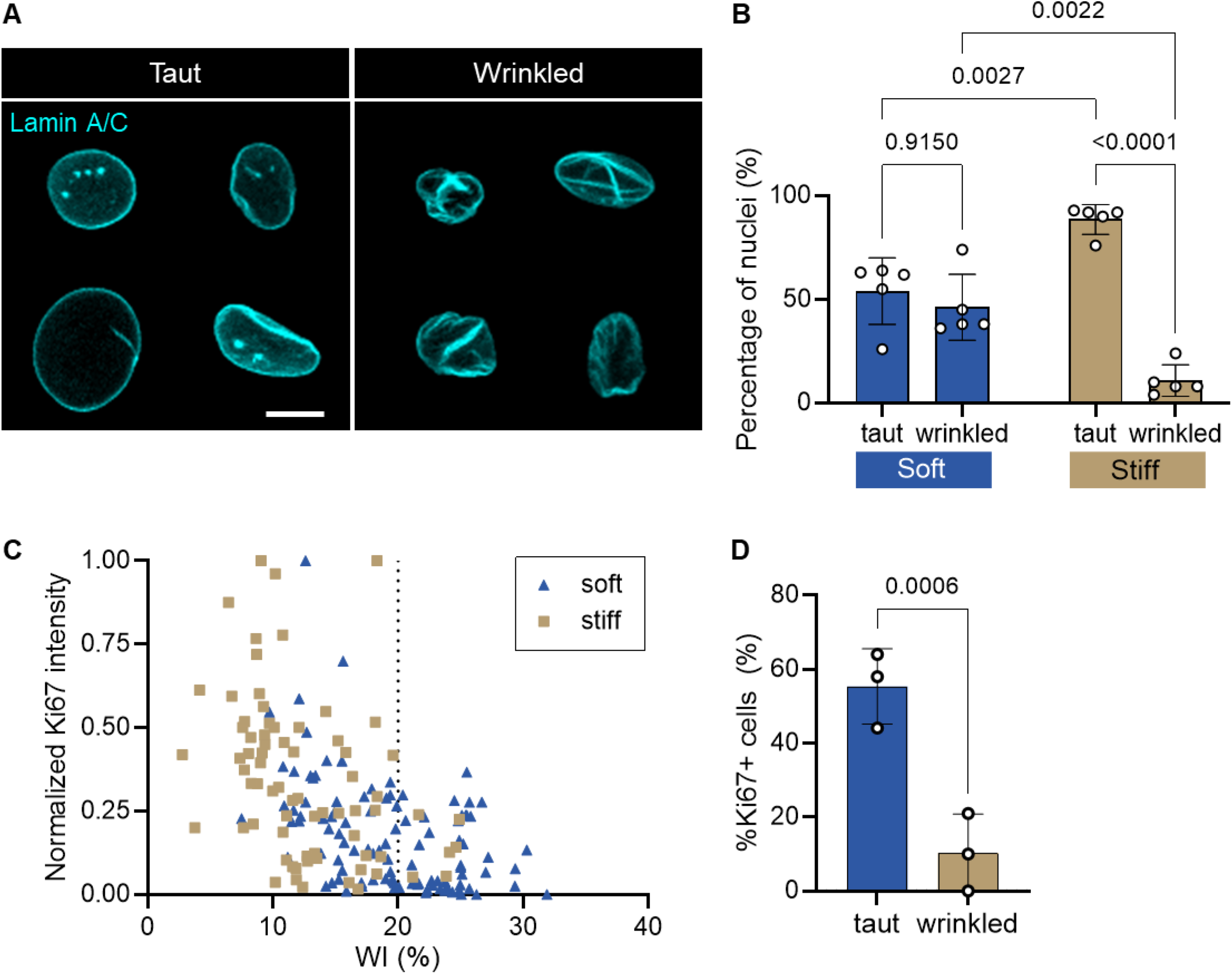
Stiffness-dependent heterogeneity in nuclear envelope wrinkling predicts Ki67 status. **A** Representative confocal images of taut (wrinkle index (WI)≤20) and wrinkled (WI>20) nuclear envelopes. Lamin A/C-cyan. Scale bar=10 μm. **B** Bar graph showing the percentages of taut and wrinkled nuclei on soft and stiff substrates. Each data point represents the mean of one independent technical replicate. Error bars report mean ± SD. n=149-121 cells per condition across N=5 independent experiments on 3 different cell lines. Statistical comparison was made by one-way ANOVA followed by a Holm-Šídák multiple comparison test **C** xy plot with x=WI (%) and y=normalized Ki67 fluorescence intensity (arbitrary unit). Each data point represents one cell. n=180 cells total across N=3 biological replicates **D** Bar graph comparing the percentage of Ki67+ nuclei in taut and wrinkled nuclei populations. n=180 cells total across N=3 biological replicates. Statistical comparison was made by a paired one-tail t-test.

We further plotted Ki67 intensity against the WI of each cell in each stiffness condition. Ki67 intensity levels were either negative, low positive (cells in active cell cycle but not S-phase), or high positive (cells in S-phase)[44]. We observed that although a taut nucleus did not always have high Ki67 intensity, a nucleus with high Ki67 intensity was always taut (WI ≤ 20%), at least in the three cell lines studied (Fig 5C). Wrinkled nuclei were four times less likely than taut nuclei to be Ki67+ at any intensity (Fig 5D).

In summary, there were higher incidences of wrinkled nuclei when myoblasts were cultured on soft substrates, and myoblasts with wrinkled nuclei did not express high levels of Ki67.

### Pharmacological manipulation of contractility modulates NE wrinkling level and myoblast proliferation

Cellular contractile force is transmitted by actin retrograde flow from adhesion sites towards the cell center, where the cytoskeleton is directly linked to the nucleus[45–47]. To investigate if low contractility on soft substrates causes NE wrinkling in myoblasts, as has been observed in mesenchymal stromal cells (MSCs)[35], we mirrored established methods to pharmacologically inhibit and activate cellular contractility using a myosin light chain kinase inhibitor (ML7) and lysophosphatidic acid (LPA), respectively. Consistent with MSCs studies, we found that ML7 treatment increased the percentage of wrinkled nuclei on stiff substrates, while LPA treatment lowered the incidence of wrinkled nuclei on soft substrates, although the effect was not as pronounced as ML7 (Fig 6A-C). Additionally, overnight treatments of ML7 decreased the proportion of Ki67+ myoblasts present on stiff culture substrates, while LPA increased Ki67+ proportion on soft substrates (Fig 6D).

**Fig 6.**
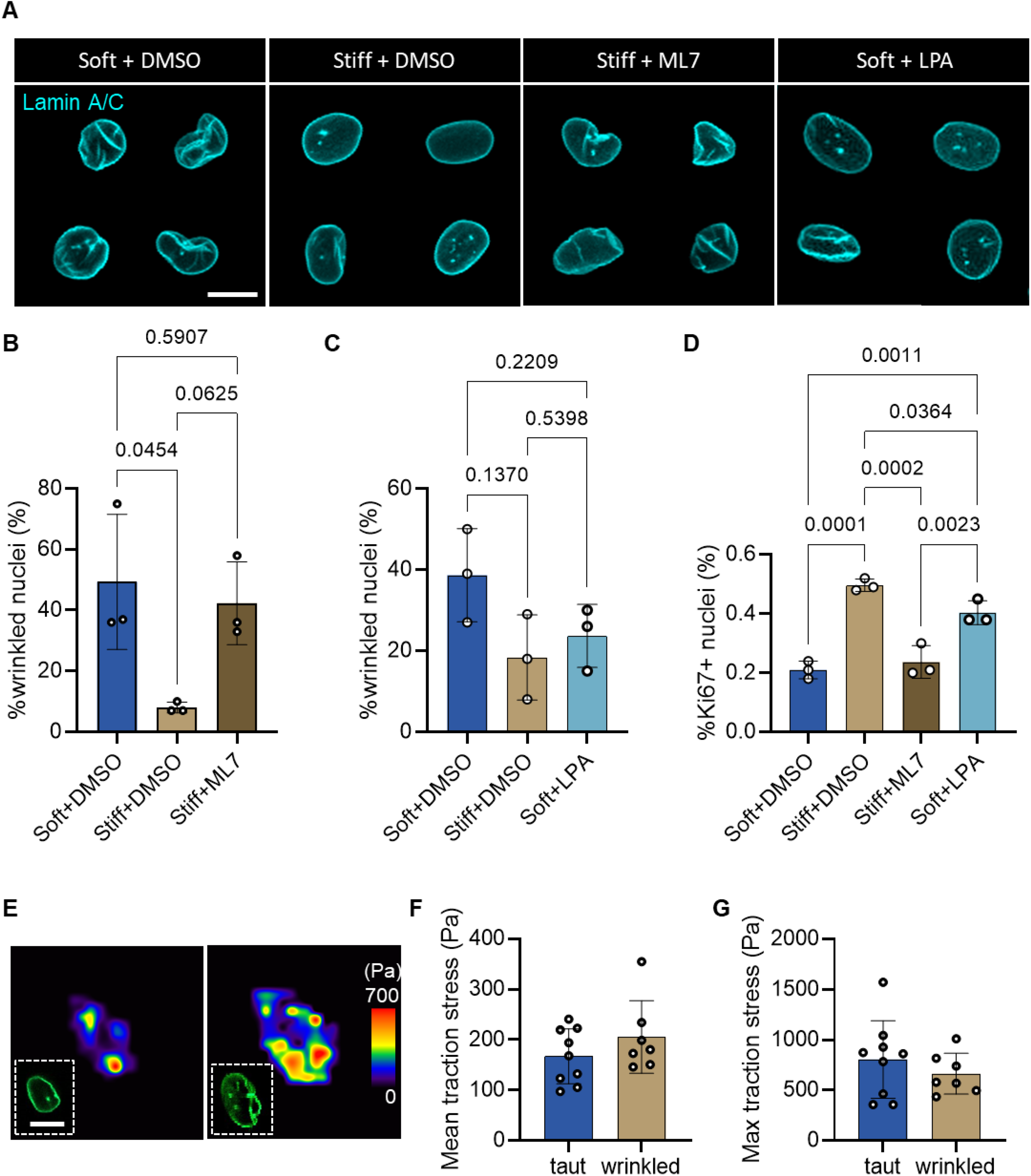
ML7 and LPA treatments alter myoblast nuclear envelope wrinkling and proliferation. **A** Representative confocal images of nuclear envelope wrinkling in myoblasts cultured on soft (1 kPa) fibronectin-tethered polyacrylamide substrates treated with either DMSO (soft+DMSO) or LPA for 2hr (soft+LPA), and in myoblasts cultured on stiff (17 kPa) fibronectin-tethered substrate treated with either DMSO (stiff+DMSO) or ML7 1hr before fixing (stiff+ML7). Lamin A/C-cyan. Scale bar=10 μm. **B** Bar graph showing the percentages of wrinkled nuclei (WI>20) in soft+DMSO, stiff+DMSO, and stiff+ML7 conditions. n=74-106 cells per condition across N=3 biological replicates. **C** Bar graph showing the percentages of wrinkled nuclei in the population of myoblasts cultured in soft+DMSO, stiff+DMSO, and soft+LPA conditions. n=60-72 cells per condition across N=3 biological replicates. **D** Bar graph showing the percentage of Ki67+ myoblasts when cells cultured on stiff and soft substrates were treated with, respectively, ML7 and LPA, both at 10 μM overnight, in comparison with DMSO controls. n=657-1060 cells across N=3 biological replicates. For all graphs, each data point represents the mean of one biological replicate. Error bars report mean ± SD. Statistical comparison was made by one-way ANOVA followed by a Holm-Šídák multiple comparison test. **E** Representative traction stress maps of a cell with a taut nucleus and a cell with a wrinkled nucleus. Scale bar = 10 µm. **F-G** Bar graph showing the mean and max traction stress of cells with taut and wrinkled nuclei. n=7-9 cells in each condition across two technical replicates of one Lamin A-GFP transduced cell line.

With these results, we hypothesized that cellular traction stresses would be lower in cells with wrinkled nuclei (WI > 20) compared to cells with taut nuclei (WI ≤ 20%). To test this hypothesis, we conducted traction force microscopy on myoblasts cultured on 17 kPa FN substrates expressing a live Lamin A reporter, the pLVX-EF1a-GFP-LaminA-IRES-Hygromycin plasmid [37]. Unexpectedly, traction stresses and strain energy values were similar between cells with taut and wrinkled nuclei (Fig 6 E-G, S4 Fig). This suggests that the relationship between cellular contractility and NE wrinkling is not straightforward. It is possible that cells with wrinkled nuclei on 1 kPa substrates had lower traction stresses than those on 17 kPa. But if low contractility was the only cause of NE wrinkling, a negative correlation between contractility and WI would have been observed among cells on the same substrate stiffness.

In summary, it appears that pharmacological manipulation of cellular contractility can alter NE wrinkling, and the absence of NE wrinkling seems to be a prerequisite for high Ki67 expression in myoblasts. However, low cellular contractility is likely not the only cause of NE wrinkling.

### Proteomic assessment of myoblast cellular processes and fate

To gain a comprehensive understanding of cellular processes following the initial responses to substrate stiffness, we profiled the proteomes of 100,000 cells cultured for 45 hours on either 1 or 17 kPa FN-tethered substrates, between which we saw the most distinct differences in cellular-level responses. We will refer to these conditions as “soft” and “stiff” substrates in subsequent discussions. 100,000 cells in each stiffness condition were split into three technical replicates. All proteomic data and analysis are provided in the supporting information.

Over 3216 protein groups were detected across the three technical replicates, and only groups with quantitative information in at least 2 out of the 3 replicates were considered. Myoblasts cultured on soft and stiff substrates had distinguishable proteomic profiles (S5A Fig). Among these filtered protein groups, 838 were determined to be differentially expressed by a p < 0.05 cut-off (S5B Fig). Using the PANTHER database to analyze differentially expressed proteins, we identified 5 enriched pathways: pentose phosphate pathway, glycolysis, integrin signaling pathway, cytoskeletal regulation by Rho GTPase, and Huntington disease (Fig 7A). The enrichment of integrin signaling and cytoskeletal regulation by Rho GTPase are expected in the context of adherent cell stiffness-sensing. Cancer cells cultured on soft environments have been reported to have reduced metabolic activities which may play a role in decreasing proliferation rate[48]. With the top two overrepresented pathways in our proteomic analysis being pentose phosphate pathway and glycolysis, we further looked at the expression of components of these metabolic pathways, and ATP synthases, for evidence of myoblasts potentially being metabolically dormant on soft substrate. Many enzymes catalyzing chemical processes in pentose phosphate pathway and glycolysis were detected to have been more abundant on soft substrates, and ATP synthase subunits expressions had mixed correlation with substrate stiffness (S6A-B Fig). Without measuring direct metabolic outputs such as ATP levels, whether there was a metabolic halt on soft substrates remained a standing question, but our proteomic assessment suggests that if there was a halt, it likely would not have been due to the lack of metabolic enzymes or ATP synthase.

**Fig 7.**
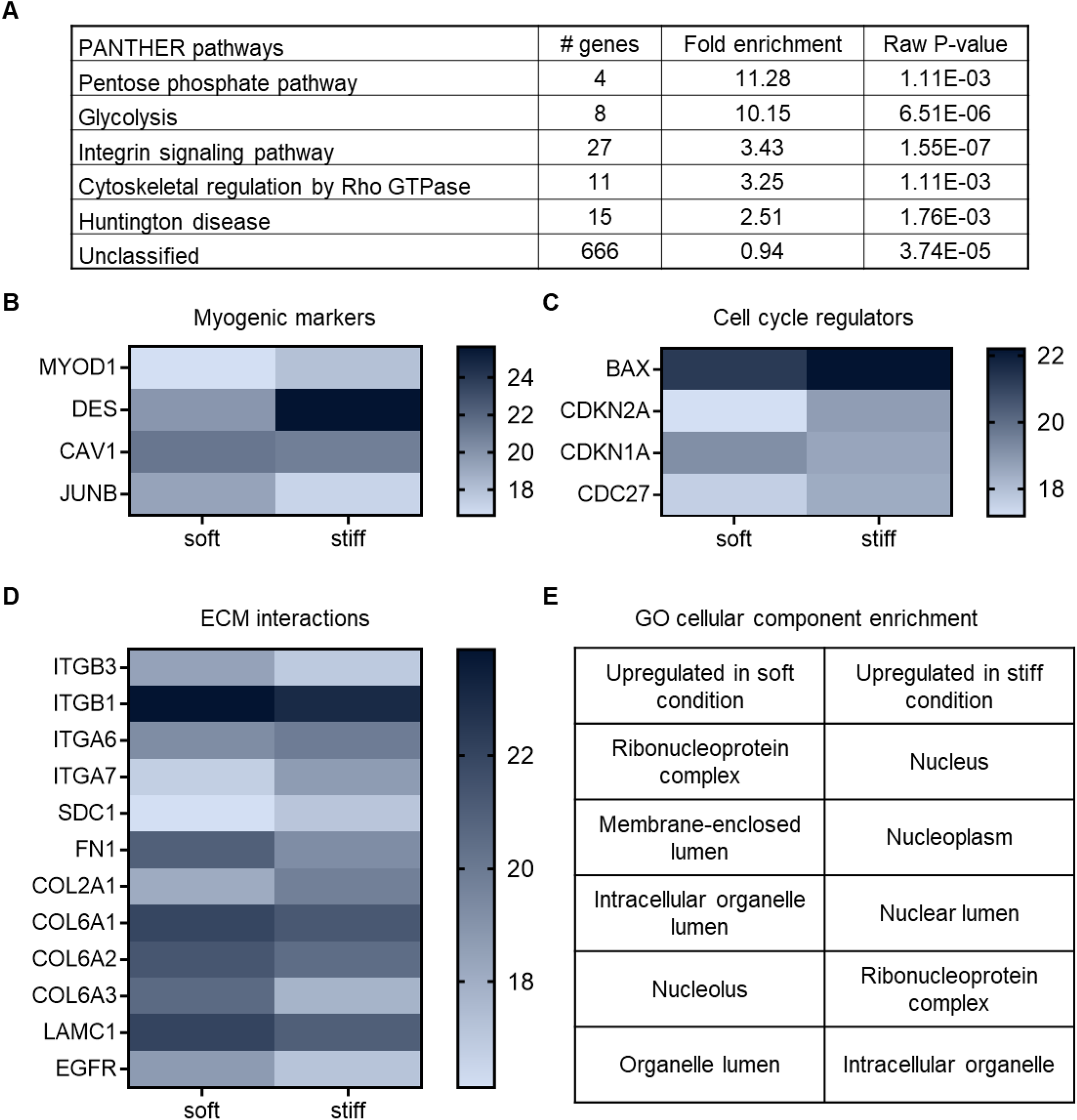
Profiling myoblast proteomes on fibronectin-tethered soft (1 kPa) and stiff (17 kPa) culture substrates at 45 hr post-seeding. **A** Overrepresented PANTHER pathways from the list of differentially expressed proteins upon comparing human myoblasts cultured on soft and stiff fibronectin-tethered polyacrylamide substrates **B** Differentially expressed myogenic-relevant markers **C.** Differentially expressed major cell cycle regulators **D** Differentially expressed proteins with direct ECM interactions **E** Table showing top five enriched cellular components from GO cellular component annotation of proteins upregulated in soft and stiff substrate conditions.

Myogenic proliferation and differentiation are regulated by a series of transcription factor activities. In vitro, myoblast fate signatures are often profiled by assessing expression of the transcription factors Pax7 and MyoD: Pax7+/MyoD-(stem-like or reserve-like cells), Pax7+/MyoD+ (proliferative cells), and Pax7-/MyoD+ (differentiating cells). We asked whether there was an increase in the proportion of either reserve-like or differentiating cells that might have corresponded to the larger portion of non-proliferative cells observed on soft culture substrates. We found that the proportions of fate signatures were similar on both stiffnesses, with most cells being Pax7+/MyoD+ (S7A-B Fig). Interestingly, the proteomics data reported an overall lower MyoD expression for myoblasts cultured on soft substrates (Fig 7B), while desmin, a myogenic commitment marker, was expressed at a higher level on stiff substrate (Fig 7B). Furthermore, JUNB, which has been previously reported to suppress myogenin expression[49], was found to express at a higher level on soft substrate (Fig 5 B). Our immunocytochemical analysis suggests that there may not have been a major divergence in the Pax7/MyoD myogenic fate landscape across the two substrate stiffness levels at the 45-hour time-point post-seeding in growth medium. However, based upon the proteomics assessment a stiff substrate at 17 kPa would favor a more differentiative phenotype while soft substrate would favor a more reserve-like phenotype. To significantly cultivate these fate transitions, we expect that other factors like culture time and medium composition would need to be redesigned.

We noticed higher expressions of ECM proteins like FN1, several COL alpha units, and LAMC1 in cells on soft substrate (Fig 7D), which could suggest that myoblasts were actively attempting to modify their environment thus not being completely inactive on soft substrate.

We also performed gene ontology (GO) overrepresentation tests on proteins upregulated in either soft or stiff condition. For these tests, we further filtered differentially expressed proteins by their fold change (stiff/soft). Only proteins with fold change less than or equal to 0.9, or more than or equal to 1.1 were considered. Among the top five enriched cellular components in either stiffness were several nuclear components (Fig 7E). LMNA expression level was found to be higher on stiff substrate. Since outside-in environmental signal transduction into cellular function converges at the level of transcription, major changes happening in the nucleus could provide important insights into how substrate stiffness is transduced into a proliferation response.

## Discussion

In this work, we characterized the short time-point, ECM ligand-specific cellular responses of primary human myoblasts cultured on mechanically tuned polyacrylamide-based hydrogels, described the substrate stiffness-specific proteomes of human myoblasts, and reported that nuclear envelope wrinkling can predict myoblast cell cycle state. Our study revealed that FN ligand tethering led to more exacerbated substrate-stiffness-dependent morphological, contractile, and proliferative responses in myoblasts compared to LAM and COL. We reported that the myoblast proteome responds to stiffness by differentially regulating metabolic, Rho cytoskeletal regulation and integrin signaling pathways, with interesting enrichment of nuclear components. Lastly, heavily wrinkled nuclear envelope morphology can predict the absence of Ki67 expression, which can be restored by LPA treatment.

The dependence of stiffness-sensing mechanics on ECM ligand is an understudied realm of myogenic cell mechanosensing. ECM ligands bind to specific integrin receptors, which have distinct molecular force transmission mechanics[26]. For example, the FN-integrin α5 bond requires the tension-dependent conformational change of FN to expose a hidden energy site, while COL can bind tension-independently to integrin α2, leading to substrate stiffness-dependent FAK activation on FN-presenting substrates but not those presenting COL[50]. The effects of ECM ligands on myogenic cell stiffness-sensing appear to vary depending on the species and passage number of myogenic cells studied and the specific experimental parameters [17,27,28]. In our experiments, we observed that relative to LAM and COL, culturing myoblasts on FN-tethered hydrogels led to more exacerbated stiffness-sensitivity in terms of cell spreading, adhesion alignment, cellular traction, and proliferation. At longer culture time and higher stiffness range, murine MuSCs proliferation have been shown to be stiffness-dependent on both LAM and RGD-presenting hydrogels, while MyoG commitment was stiffness-dependent on RGD and not LAM[17]. Confounding factors to PA hydrogel Young’s modulus, such as porosity and ligand tethering density could potentially add to result variability[51,52].

The idea that ECM composition could potentially alter myogenic cells mechano-sensitivity is important in the context of in vivo niche ECM remodeling during regeneration, aging and disease. MuSCs niche factors were reported to actively suppress activation pathways to maintain MuSCs quiescence[53,54]. As for LAM, past studies have shown that laminin-2/4 deficiency leads to the loss of integrin α7β1[55], which in turn could disrupt quiescence maintenance[56]. We can therefore infer that the SC connection to LAM via integrin α7β1 is important in preserving SC quiescence, but the exact mechanisms are unknown. In vitro studies on myoblasts, in agreement with other cell types, showed that LAM preferentially activates Rac and suppresses RhoA[57]. In contrast with FN, LAM promotes motile cells with diffused stress fibers and poor vinculin-containing adhesion assembly, characteristic of a high Rac/low RhoA phenotype[58]. These observations are exciting in the context of the Rac-to-Rho switch occurring during MuSCs activation[59]. Rho-regulated cytoskeletal remodeling is critical in cell tension sensing, including matrix stiffness. It is therefore enticing to explore the possibility that LAM actively suppresses MuSCs mechanosensing, while FN enrichment in the injured niche increases MuSCs sensitivity to injury sensation and facilitates the quiescent to proliferative transition, and furthermore, how age- and disease-related irregular ECM deposition impair MuSCs sensitivity to niche remodeling.

MuSCs are functionally heterogeneous[60], as are their downstream progenitors. Our results suggested that relatively stiff substrates, tethered with FN, decrease myoblast cell cycle heterogeneity, cultivating a primarily proliferative population. While we observed that soft substrates tethered with FN increased the incidence of Ki67-myoblasts, we did not assess whether the cell cycle itself was impeded. Regarding the split between Ki67+ and Ki67-myoblasts on the same substrate stiffness, our data suggested that perhaps one of the differences between these subpopulations is their contractility, since high Ki67 expression was only observed in the absence of NE wrinkling, a state typically characterized by high cellular contractility[35]. To assess the correlation between cell contractility and NE wrinkling level, we performed traction force microscopy on cells expressing a live Lamin A reporter. To our surprise, cellular traction stress and strain energy were independent of NE wrinkling index, despite pharmacological manipulation of contractility resulting in expected NE wrinkling and Ki67 expression responses. There are a few possible explanations. First, pharmacological manipulation experiments were done on two substrate elasticities while traction force microscopy on Lamin A-GFP expressing cells were conducted exclusively on 17 kPa substrates, so it is possible that cells with wrinkled nuclei on 1 kPa had lower traction stresses than cells with wrinkled nuclei on 17 kPa. Second, our proteomic analysis detected a decrease in Lamin A/C on soft substrate, which could potentially take part in increasing NE wrinkling[61,62]. Third, we did not assess nuclear-actin connectivity. If there is a disruption in tension propagation from the cytoskeleton to the nucleus, nuclear tension would be low regardless of cytoskeletal contractility. For example, severing the Linker of Nucleoskeleton and Cytoskeleton (LINC) complex has been shown to induce similar nuclear wrinkling phenotypes[35,63].

Whether and how exactly NE wrinkling could suppress proliferation are questions whose pursuit would require the manipulation of NE wrinkling without altering cellular signaling, which is a limitation of the pharmacological strategies that we and others use to manipulate contractility. LPA activates a range of signaling pathways including Ras and Rho/ROCK GTPases which are cell cycle regulators[64–66]. Alternatively, low nuclear tension reduces nucleocytoplasmic exchange of signaling molecules like YAP, cyclin B1, and MyoD by potentially deforming nuclear pore complexes[67–69]. Cosgrove et al. reported a correlation between NE wrinkling and YAP/TAZ nuclear localization in mesenchymal stem cells [35]. In our system, bulk proteomic profiling on soft and stiff substrates did not detect differential YAP target proteins between soft and stiff substrates, suggesting that a YAP/TAZ localization mechanism may not underlie our results.

In conclusion, our work emphasized the integrated control of substrate stiffness and ECM composition on myoblast contractility and proliferation, which are heterogeneous phenotypes intimately linked to the NE morphology. The impact of our work extends to biomaterial design applications and implicates fundamental regulation of human myogenic cell mechanosensing.

## Supporting information

Compiled Raw Data

Proteomics Data

## Acknowledgements

We would like to thank Dr. Qingzong Tseng for advice on the implementation of his TFM ImageJ macros and Dr. Louise Moyle for thoughtful discussions. This work was funded by a Natural Sciences and Engineering Research Council grant to P.M.G. (RGPIN-2019-07144), a Human Frontiers Science Program (RGP0018/2017) to P.M.G. and T.B., a Deutsche Forschungsgemeinschaft (DFG, BE 6270/2-1) to TB, a Connaught International Scholarship to J.N., and a Canada Research Chair in Endogenous Repair to P.M.G. (#950-231201).

## Author contributions

Conceptualization: J.N., P.M.G.; methodology: J.N., L.W., M.B.; investigation: J.N., L.W., W.L., Y.H., N.G., C.C.M.; validation and formal analysis: J.N., W.L., L.W., Y.H., N.G.; visualization: J.N., muscle tissue collection and pre-processing: H.A., H.G., R.K.; writing-original draft and writing-review & editing: J.N., P.M.G.; supervision and funding acquisition: T.B., P.M.G. All authors approved the final manuscript.

## Conflict of Interest

The authors have no conflict of interests that might be perceived to influence the results or discussion reported in this paper.

**S1 Fig. Log-normal cell spreading area distribution A** Frequency distribution of raw, unadjusted myoblast cell spreading areas on 1 kPa fibronectin-tethered polyacrylamide substrates **B** Frequency distribution of the natural log of the same cell spreading area data as in A. **C, D, E** Graphs showing the natural log of myoblast spreading area before normalizing to 9 kPa condition. Each data point represents the mean of one biological replicate. n=395-920 cells per condition across N=3 biological replicates. Error bars report mean ± SD.

**S2 Fig. Viability analysis of myoblasts cultured on fibronectin-tethered polyacrylamide substrates.** Representative images of human myoblasts cultured for 16 hours on 1, 3, or 17 kPa fibronectin-tethered polyacrylamide substrates and stained for Hoechst (yellow), propidium iodide (PI; magenta) and Calcein (green). Scale bar=500 μm.

**S3 Fig. Strain energy of myoblasts across substrate stiffnesses and ECM culture conditions.** Bar graph showing strain energy of myoblasts cultured across substrate stiffness and ECM conditions. n=19-24 cells per condition across N=3 biological replicates. Each data point represents the mean of one biological replicate. Error bars report mean ± SD. Statistical comparisons were made by two-way ANOVA followed by a Holm-Šídák multiple comparison test.

**S4 Fig. Strain energy of myoblasts with taut vs. wrinkled nuclei.** Bar graph showing strain energy of myoblasts having taut (wrinkle index ≤ 20) vs. wrinkled (wrinkle index > 20) nuclei. n=16 cells in total across 2 technical replicates. Each data point represents one cell. Error bars report mean ± SD. Statistical comparisons were made by a Student’s t-test.

**S5 Fig. Proteomic profiling of myoblasts cultured on fibronectin-tethered soft (1 kPa) and stiff (17 kPa) substrates A** Intensity heat map of all protein groups detected in myoblasts cultured on soft and stiff substrate after filtering by the following criteria: at least 2 of the 3 technical replicate runs have quantitative information; and more than 2100 proteins were quantified **B** Volcano plot showing relative distributions of upregulated proteins in terms of -Log P-value and expression difference (soft-stiff) of all detected proteins. Proteomic analysis was done on one cell-line with 100,000 cells in each stiffness condition, and split into 3 technical replicates.

**S6 Fig. Expression of metabolic pathway components on soft (1 kPa) and stiff (17 kPa) substrate. A** Heat map showing protein levels of metabolic enzymes involved in glycolysis and pentose phosphate pathway in soft and stiff substrates **B** Heat map showing protein levels of ATP synthase subunits on soft and stiff substrates.

**S7 Fig. Pax7/MyoD analysis of myoblasts cultured on fibronectin-tethered soft (1 kPa) and stiff (17 kPa) substrate A** Representative confocal images of myogenic transcription factor immunostaining. Hoechst-grey, Pax7-cyan, MyoD-yellow. Scalebar=20um. **B** Bar graph showing the percentage of cells in each fate category (Pax7-/MyoD+, Pax7+/MyoD+, Pax7+/MyoD-, Pax7-/MyoD-) on soft and stiff substrate. n=435-469 cells across N=3 biological replicates.

