## Supplementary figures and images for "Culture substrate stiffness impacts human myoblast contractility-dependent proliferation and nuclear envelope wrinkling"

### Supplementary Figures

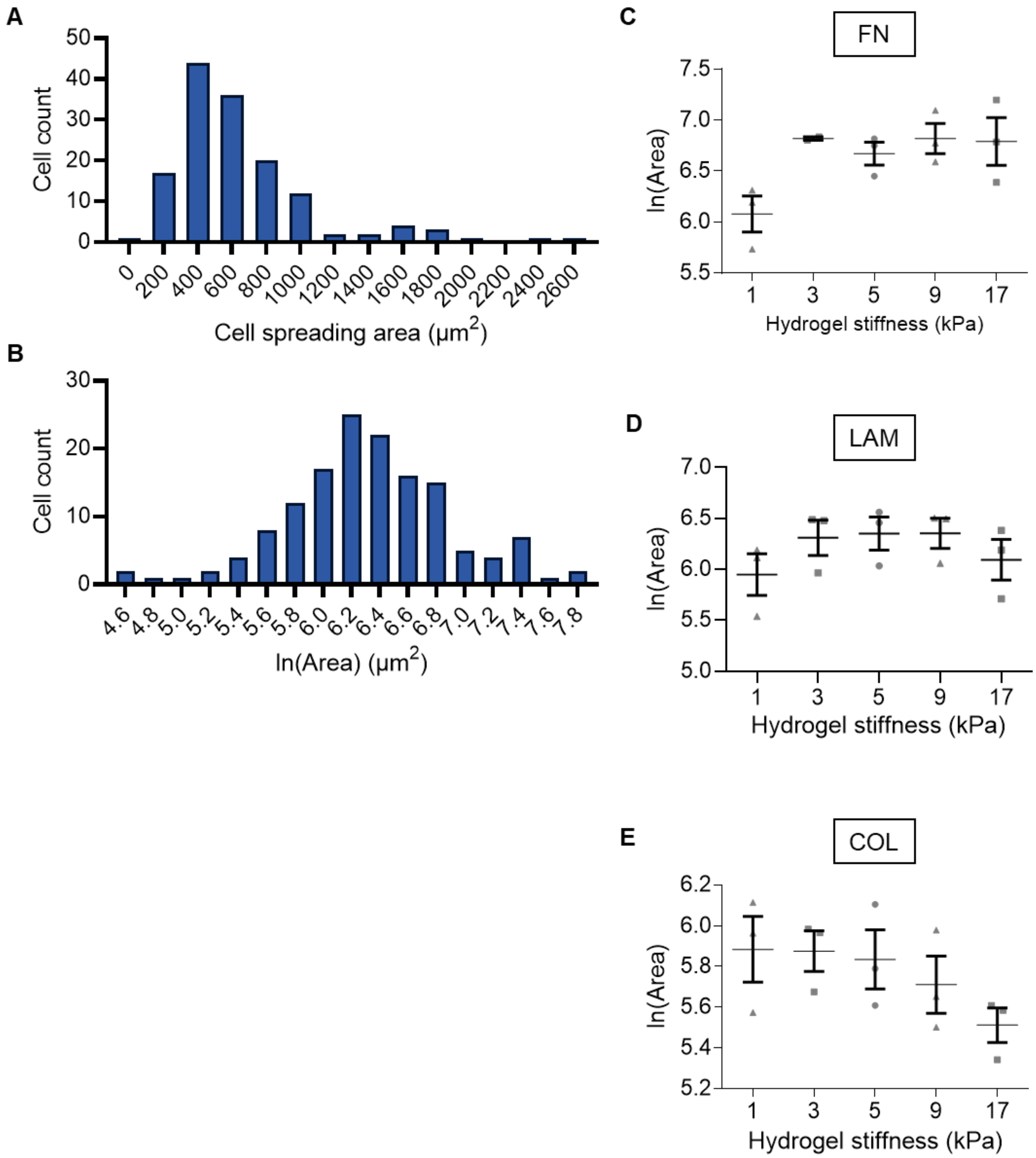

**S2 Fig.**

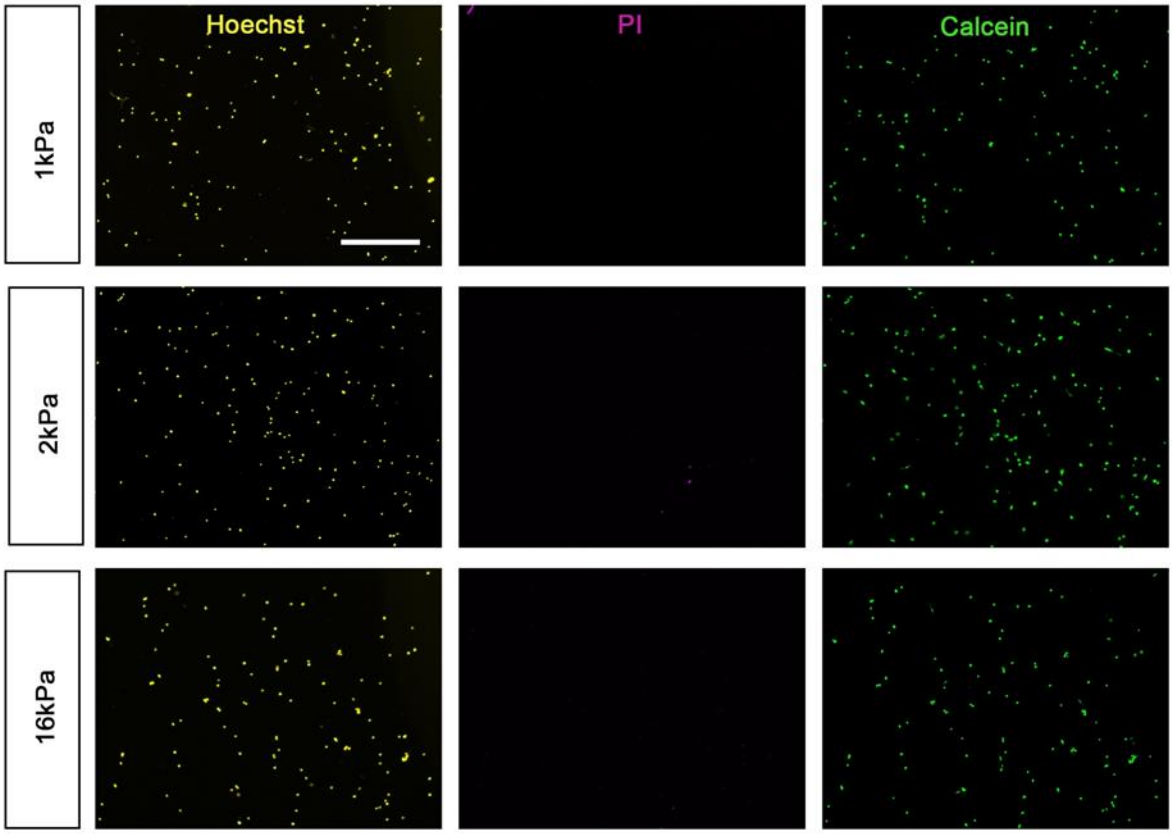

#### S3 Fig.

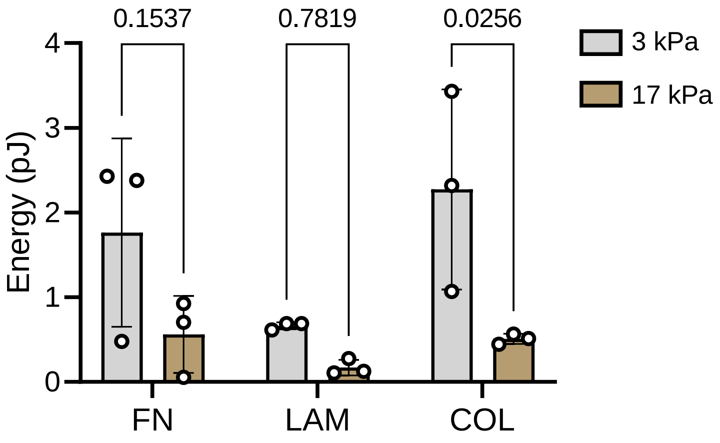

**S4 Fig.**

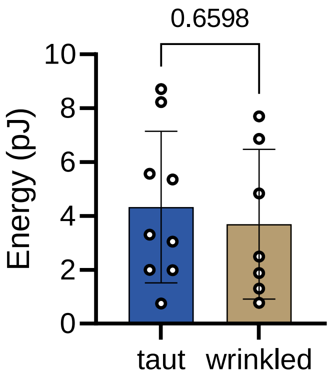

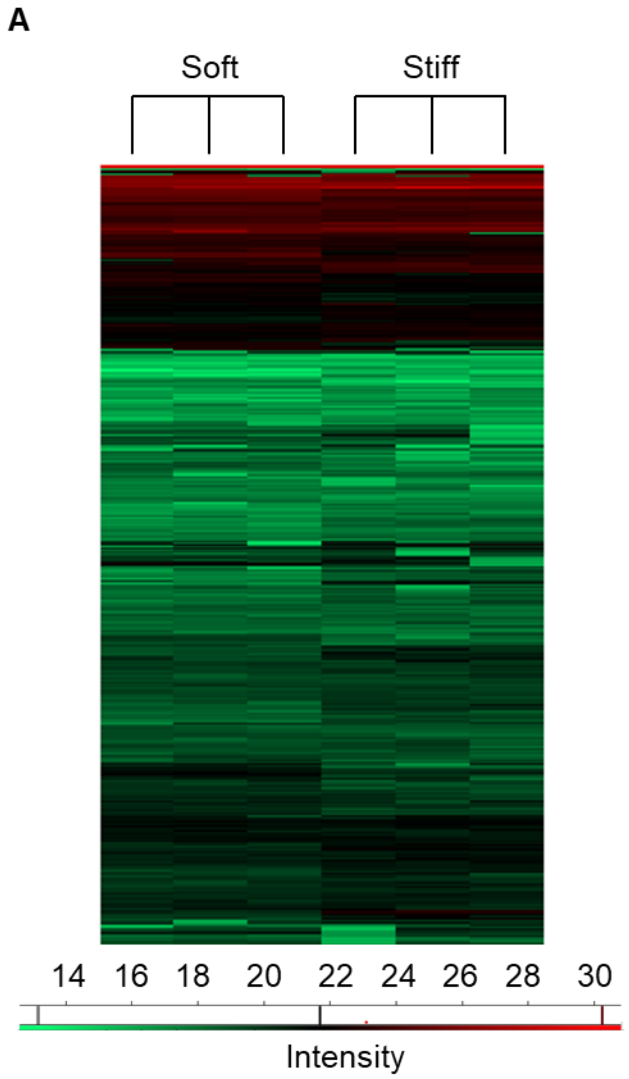

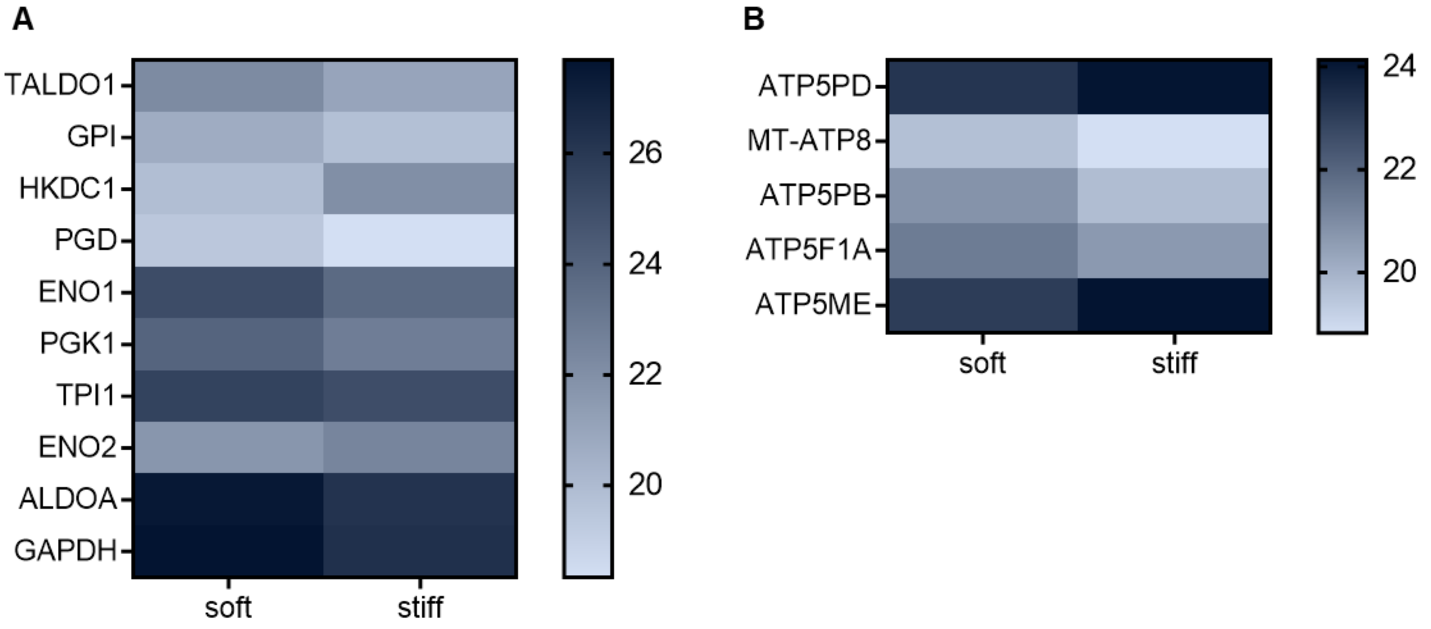

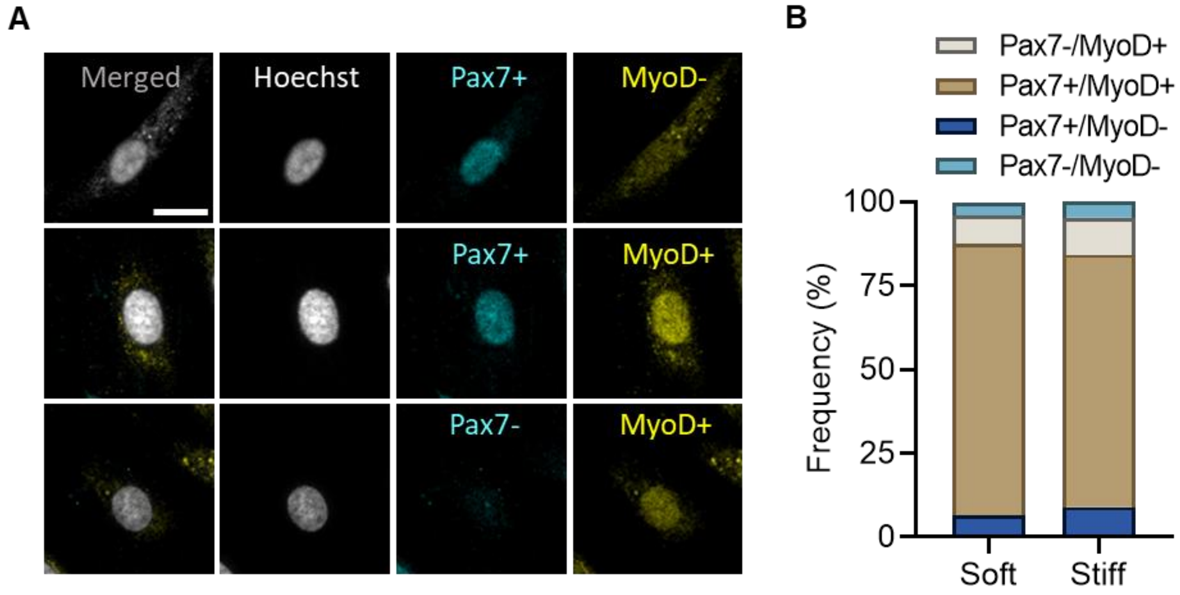
